# Recycling of Polymerase Chain Reaction (PCR) kits

**DOI:** 10.1101/2022.04.08.487615

**Authors:** Weina Liu, Francesco Stellacci

## Abstract

During the outbreak of the SARS-CoV-2 pandemic, PCR (polymerase chain reaction) kits have been used as a common diagnosing method, with daily worldwide usage in the millions. It is well known that at the beginning of the pandemic there was a shortage of PCR kits. So far, the ecosystem of PCR kit is linear use, that is kits are produced, used one-time, and disposed in biolab wastes. Here we show that, to mitigate the risk of future shortages, it is possible to envision recyclable PCR kits, based on a more sustainable use of nucleic acid resources. A PCR kit is mainly composed of primers, nucleotides, and enzymes. In the case of a positive test, the free nucleotides are polymerized onto the primers to form longer DNA strands. Our approach depolymerizes such strands keeping the primers and regenerating the nucleotides, *i*.*e*., returning the nucleic acid materials to the original state. The polymerized long DNA strands are hydrolyzed into nucleotides monophosphates that are then phosphorylated in triphosphates using a method that is a development of a recently published one. We used oligonucleotides with 3’-terminal phosphorothioate (PS) backbone modification as non-hydrolysable PCR primers, so to undergo the recycling process unchanged. We have successfully recycled both PCR primers (∼65% yield for 4-PS modification, and ∼40% yield for 2-PS modification) and nucleotides (∼75% yield). We demonstrate that the method allows for direct re-use of the PCR kits. We also show that the recycled primers can be isolated and then added to end point or quantitative PCR. This recycling approach provides a new path for circularly reusing PCR nucleic acids.

## INTRODUCTION

A ‘circular economy’ would change the economic logic by turning linear used old-goods into as-new resources with the aim to form closing loops in industrial ecosystem, minimize waste, and lessen the need to make originals from scratch.^1^ For polymers, de-polymerisation is required to produce smaller molecules (often the starting monomers) that can be re-used to produce new polymers or new molecules.^2,3^ Our group has recently reported a recycling approach for sequence-defined polymers (SDP) that we call NaCRe (nature-inspired circular-recycling system), we first implemented it onto proteins^4^ and more recently onto DNA.^5^ Briefly, NaCRe’s concept is that in theory a random mixture of SDPs can be depolymerized into the constituent monomers that in turn are re-assembled in a new polymers that can potentially differ from any of the starting ones. This has been shown for proteins (SDPs made of amino acids) and for DNA SDPs made of nucleotides. We showed that commercially available calf-DNA can be hydrolyzed into monophosphate nucleotides (dNMPs) that then must be enzymatically converted into their triphosphate counterparts (dNTPs) in order to be further used to produce DNA strand by PCR. Indeed, we proved that the latter ones can be used as reagents in PCR to produce new DNA strands not related to the original ones. NaCRe for DNA could be useful to render more sustainable applications that use nucleotides as reagents such as molecular diagnosing, and sequencing,^6,7^ as well as applications where DNA is used as materials such as DNA origami, DNA hydrogel, and DNA for data storage.^8–10^ Yet, at present arguably the main technological use of DNA is in PCR kits. NaCRe as such cannot be used for PCR kits because, as described, it would hydrolyze (depolymerize) the main and most expensive nucleic acid reagent, the primers. Here we show that we can combine DNA NaCRe system with primers that are suitably terminally protected to achieve a system that, upon hydrolyzation, regenerates both the original primers and the nucleotides.

Since the worldwide outbreak of the SARS-CoV-2 pandemic, massive PCR testing is required in many countries for diagnosing and screening.^11,12^ At the end of 2021, the worldwide cumulative covid test (PCR and antigen test) reached ∼4 billion.^12^ PCR testing kits are single-use and end up in the waste after testing. A large amount of biolab waste is accumulated through this massive, daily repeated PCR test. There are many methods established for the *de novo* synthesis, conjugation, and amplification of DNA with varied lengths and functions. They include (1) solid-phase phosphoramidite synthesis of oligonucleotides with customized sequences,^13^ (2) enzymatical synthesis of template-independent oligonucleotides by polymerase-nucleotide conjugates,^14^ (3) DNA printing for the synthesis of large libraries of DNA strands,^15^ (4) phagemid assisted scalable and efficient production of staple DNA for the construction of DNA origami,^16^ (5) Gibson assembly for the high throughput, end-linkage of short DNA for the construction of genome DNA.^17^ To the best of our knowledge, the de-polymerization and recycling of DNA materials had not been reported before our recent publication.^5^

A PCR kit is basically a DNA polymerization and amplification tool. Briefly, PCR is a polymerase catalyzed, templated, temperature-controlled, precision DNA polymerization process. In a PCR kit, there are primers, nucleotides, and enzymes. The function of primers (oligonucleotides) is for sequential recognition and complementary binding to template DNA (e.g., the representative sequences of viral genomes), and initiation of the chain elongation for DNA polymerization. PCR polymerization proceeds in the 5’ to 3’ direction.^18^ Enzymes drive the polymerization consuming the nucleotides (the monomers in this reaction). The PCR waste is a mixture of positive and negative PCR products. In case of a positive test, the final products are many copies of the targeted DNA strand, while a negative test has the oligonucleotides in the original state. To recycle PCR kits, a controlled de-polymerization method is required. To be successful, this depolymerization must proceed in the opposite direction when compared to the PCR DNA polymerization (that is in the 3’ to 5’ direction) in order to preserve suitably modified primers. In other words, the depolymerization should stop at the primer leaving the primer intact and ready to be reused. A primer modification with the function of hydrolyzation-stopper is required. PCR primers are oligonucleotides synthesized by solid-phase chemistry. During the synthesis, chemical modifications of the nucleobases, sugar bond, as well as inter-nucleotide phosphodiester bond can be readily introduced into the primers.^19,20^ In this paper we used a phosphorothioate (PS^21^) modification at the 3’-terminal developed as a nuclease-resistance backbone-modification^20,22,23^ for oligonucleotides drugs as a non-hydrolizable PCR primer. The effective combination of NaCRe^5^ together with PS-modified primers allowed us to efficiently recycle both primers and nucleotides from PCR kits with a process that we illustrate in Scheme 1. This DNA recycling approach brings a new perspective to the design of PCR kits.

## RESULTS AND DISCUSSION

### PS-Modified DNA

To establish our method for recycling PCR kits, we first needed to determine whether the hydrolysis resistant oligonucleotides we chose could be used as primes in a PCR experiment. Forward and reverse primers were designed with a 4-PS tail in their 3’-terminal (Scheme 1). The PS-modified oligonucleotides were used as PCR primers for the amplification of DNA template with luciferase sequence (luc, amplification length 1653 base pair^24^). The primers without PS-modification were applied for PCR as control to compare the polymerization efficiency. The PCR amplification products were purified using a standard PCR extraction kit and quantified using Nanodrop.^25^ We found only a slight decrease of PCR yield by using the PS-modified primers (Figure S1a), showing that the function of primers binding to their DNA targets and the polymerase catalyzed chain extension was not strongly affected by PS-modification.

To test the resistance of such backbone modified primers to hydrolysis we used a method we recently developed. Indeed, we established a one-pot DNA hydrolysis method^5^ to produce dNMPs in good yields. Fluorescence dye-labeled primers (with or without PS-modification) were incubated together with nucleases to evaluate their nuclease tolerance. The reaction mixture was loaded to a 2% agarose gel. Very good hydrolysis resistance capacity of PS-modified primers was observed (Figure S1b). The PS-modified primers can tolerate more than 24h nuclease hydrolysis. In comparison, the primers without PS modification were hydrolyzed after 2h. The result shows that the 4PS-modification provides excellent protection against nucleases-induced hydrolysis of PCR primers. Therefore, the PS-modified oligonucleotides can be used as recyclable primers for PCR.

### Recycling of PCR kit

We then used such PS-modified primers in a PCR kit that lead to successful DNA elongation (Scheme 1, step 1). The whole process of DNA polymerization (PCR) and de-polymerization (recycling) was monitored by HPLC (Figure S2). After PCR amplification, about two thirds of dNTPs were consumed (Figure S2d). In Figure S2a (lines 1 and 2) we show retention time plots obtained using HPLC (high performance liquid chromatography). In such plots when comparing a fresh PCR kit to a used one, we find that as expected dNTPs are consumed due to the polymerization. We also find new peaks of nucleotides diphosphates (dNDPs, Figure S2a, line2). We believe that these dNDPs derive from dNTPs due to their at least partial hydrolysis induced by heating in PCR condition. In order to show that the overall product mixture generated by the PCR analysis is suitable for NaCRe recycling, we applied the second NaCRe steps to it (see Scheme 1, step 2), that is we overall mixture was hydrolyzed to obtain dNMPs. In this step, we modified the published protocol of hydrolysis in order be more efficient in hydrolyzing all DNA templates and the PCR product that would be precent in a standard positive PCR kit. Two restriction enzymes (BanI and BstYI) with multiple cleavage sites of the amplified luc sequences were added to the PCR product so that the luc sequence can be fragmented into shorter pieces. Next, the DNA fragments were hydrolyzed by a mixture of exonucleases to release dNMPs, as in the established method.^5^ After the enzymatical depolymerization process, the amount of dNMPs in the recycled PCR kit was highly increased (Figure S2a, line 3). Further, the released dNMPs were phosphorylated to generate dNTPs (Scheme 1, step 3), as in the established one-pot phosphorylation method.^5^ Briefly, the phosphorylation reaction was catalyzed by T4 NMP Kinase and E. coli S30 extract (rich in phosphotransferase), with acetyl-phosphate/ATP as dual-phosphate-donors. After phosphorylation, the dNMPs peaks almost disappeared, and the intensity of dNTPs peaks increased (Figure S2a, line 4). Overall, through this hydrolysis-phosphorylation process, the dNTPs in the PCR substrate were regenerated from PCR products/waste.

The concertation of dNTPs from fresh and re-generated PCR kit was quantified (Figure 1a, with calibration curves of dNTPs in Figure S2b). After all the steps of dNTPs recycling, the reaction mixture was diluted by 1.47 times with the average concentration for each dNTP of ∼100 μM (Figure S2c). The average recycling efficiency was calculated to be 74.5 ± 2% (Figure 1b), which is relatively high, showing the possibility to recycle monomeric nucleotides materials in PCR kits. The phosphorylation yield was relatively good (on average 93 ± 2%, Figure S2d), which is similar to our reported phosphorylation yield of dNMPs.^5^ In the re-generated PCR kit, dNDPs residue was 5.94 ± 0.04%, ATP/ADP residue was 12.23 ± 0.02%, and dNTPs content was 81.81 ± 0.02% (Figure 1c). We notice that a fraction of the dNTPs was not fully hydrolyzed and retained as 3mer-6mer oligonucleotides in the recycled PCR kit (Figure S3).

**Figure 1.**
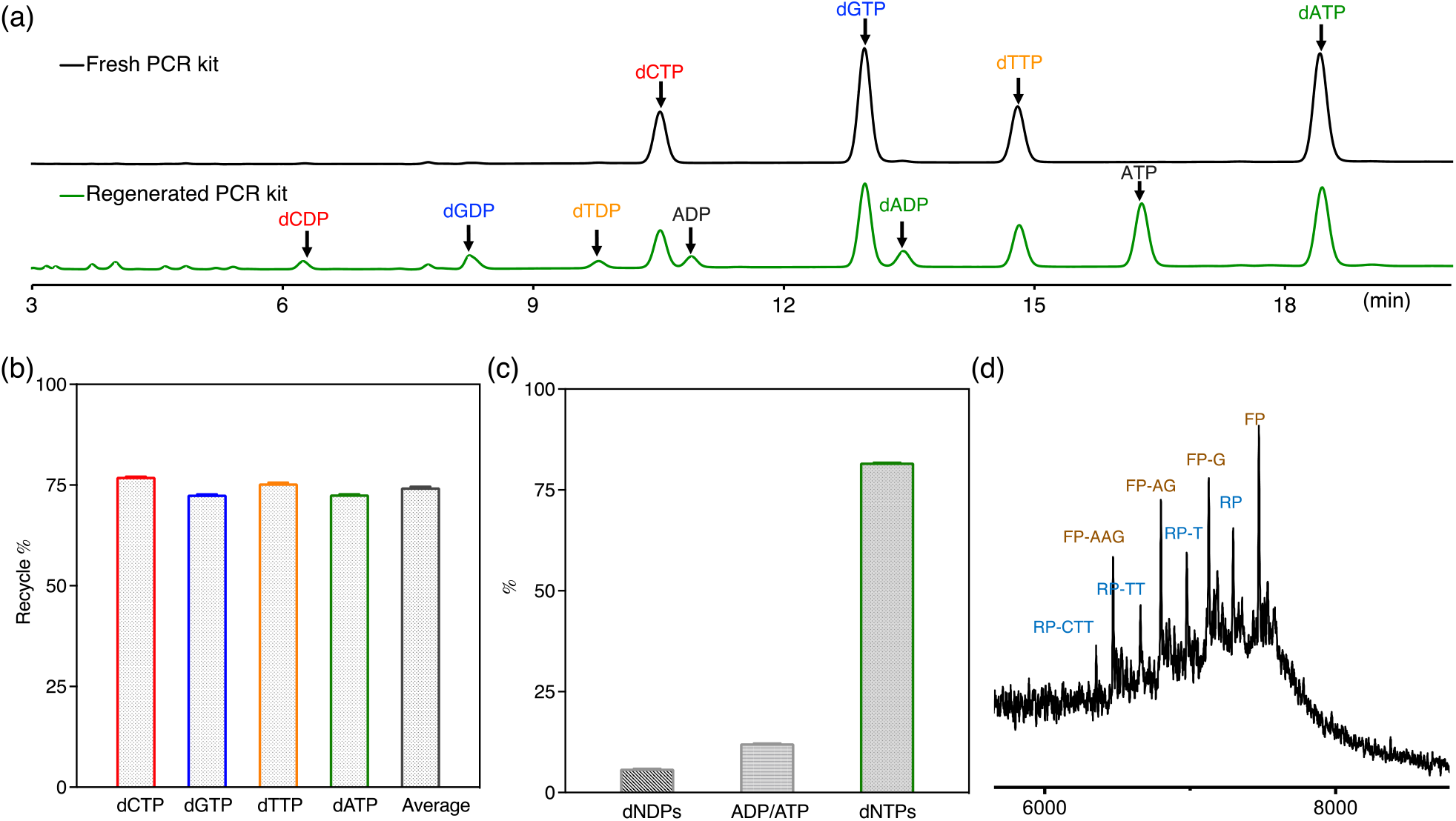
HPLC retention time of dNTPs standards from fresh PCR kit, and the products of phosphorylation mixture from the regenerated PCR substrate. (b) Recycling yield of dNTPs in the regenerated PCR substrate, dC 77.04 ± 0.05%, dG 72.64 ± 0.02%, dT 75.42 ± 0.19%, Da 72.65 ± 0.04%, in average 74.44 ± 2.01%, calibration curves of dNTPs and dNDPs see Figure S2b and 2c. (c) Ratio of all the nucleotides components in the regenerated PCR substrate, dNDPs 5.94 ± 0.04%, ADP/ATP 12.23 ± 0.02%, dNTPs 81.81 ± 0.02%. (d) Mass spectrum of recycled primers-PS with 0-3mer terminal nucleotides were hydrolyzed.

We then evaluated whether the PS-modified primers can be preserved through this enzymatic hydrolysis and recycling process. We isolated the primers using an Oligo extraction kit and the molecular mass of the extracted primers was measured by MALDI-TOF (Matrix-assisted laser desorption/ionization-time-of-flight mass spectrometer). A mixture of recycled primers with fully preserved and shortened length (loss of 1, 2, or 3 terminal nucleotides) was observed (Figure 2d), showing the 4-PS tail could effectively protect primers from the enzymatical hydrolysis step. Unexpectedly, we found that the enzymatic hydrolysis was ‘slowed down’ instead of totally ‘stopped’ by the 4-PS tail. For the preserved primers with 0 to 3 terminal nucleotides lost, the enzymatic hydrolysis was much slower, yet the remaining primers could be recycled. These residues do not account for the total mass of the initial primers. The combination of these observation leads us to believe that once all the 4PS modified nucleotide are hydrolysed, the primers quickly degrade. A possible reason for the incomplete protection to hydrolysis could be the chiral configuration of phosphorothioate backbone modification, that the *S*-PS can tolerate enzymatical hydrolysis better than *R*-PS.^26^ As the 4PS-modified primers were achieved by solid-phase synthesis, it is difficult to determine the stereo-selectivity for the PS modification at the current stage. There are ‘stereopure’ chiral phosphorothioate-modified primers reported,^27^ which could potentially perform better unders hydrolysis. Overall, the recycling yield of primers was found to be 60.85 ± 0.01 % (mass ratio) and 64.38 ± 0.01% (molar ratio, calculated from the average mass of recycled primers), which is still quite promising. We should state that due to the limited number of primers (∼ 0.7 μg) there were significant losses of material in the extraction process hence the yield we mention are lower estimates, for example the vendor claims a 90% recovery we believe ours’ recovery was lower. We have not accounted for the recovery rate when calculating the yield.

**Figure 2.**
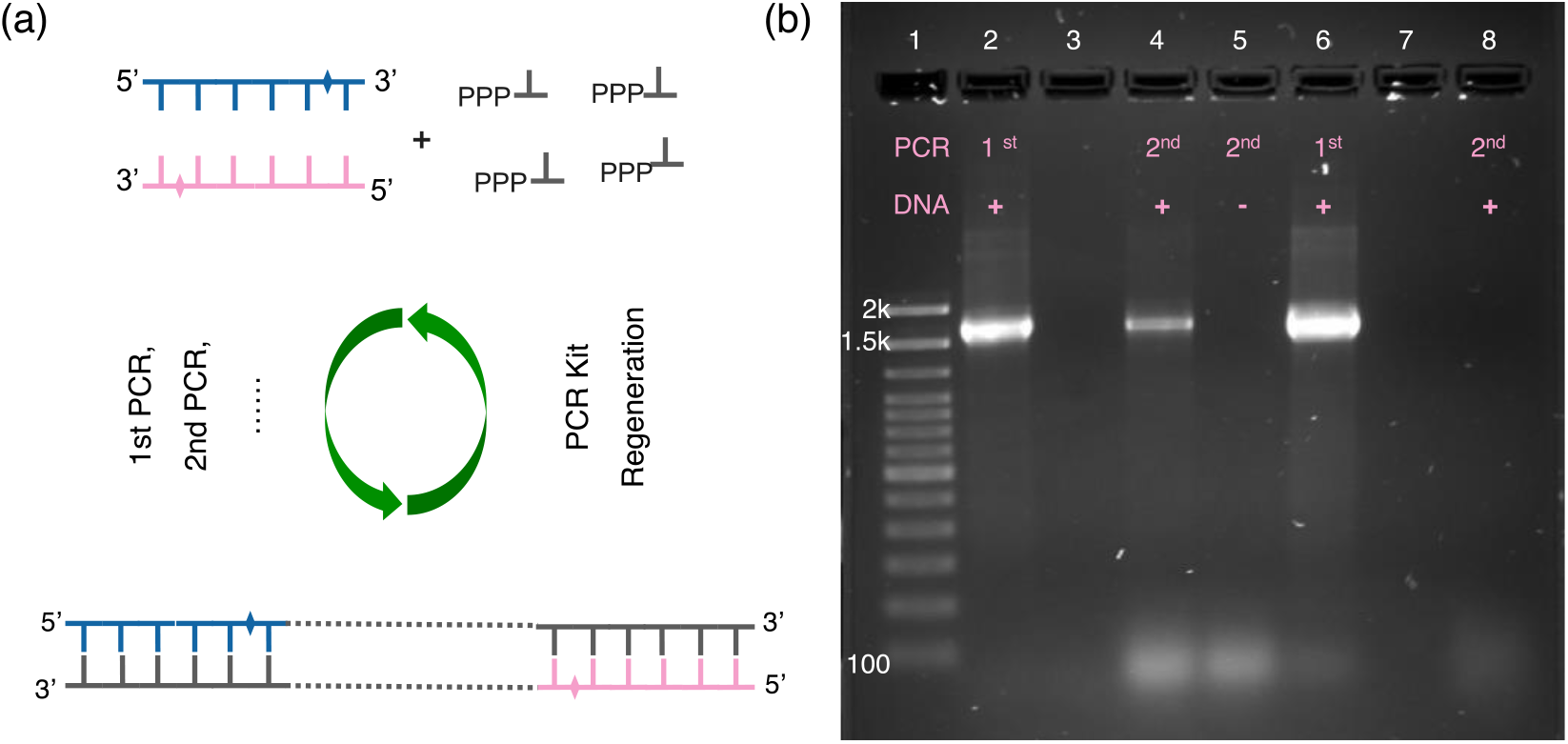
(a) A simplified scheme showing the circularly recycle and re-used PCR kit. (b) Agarose gel of PCR amplification product from fresh (1st) and regenerated (2nd) PCR kit: lane 1, ladder; lane2, 1st PCR product by using primers-PS; lane 3, hydrolyzed 1st PCR product; lane 4, 2nd PCR product with DNA template added, lane 5, 2nd PCR product without DNA template added; lane 6, 1st PCR product by using primers without PS modification; lane 7, hydrolyzed 1st PCR product; lane 8, 2nd PCR product with DNA template added.

### Reused PCR kit

In an ideal situation we would like to establish a method, to regenerate and circularly reuse PCR kits simply adding polymerization and de-polymerization enzymes when needed (Figure 2a). Therefore, we tested the possibility of reusing the regenerated PCR substrate for a second round of PCR amplification. Only new DNA templates and polymerase were added to the regenerated PCR kit for the second round of PCR. The PCR amplification products (first and second, with or without adding DNA template) and hydrolyzed PCR product were loaded to an agarose gel. In Figure 2b, we show the result of a positive PCR process that in a gel has a clear band due to the elongation of the primers (lane 2). In lane 3, the hydrolysed PCR material is shown, that band of lane 2 cannot be seen revealing successful de-polymerization of the PCR product. In lane 4 we placed the result of a PCR process performed with a fully recycled kit. As expected, the band due to primer elongation can be seen again in this lane. The intensity of the band in lane 4 is weaker than in lane 2, there are two reasons for this. First there was a dilution when going for the original to the regenerated kit (overall 1.47x dilution), which led to a decreased DNA polymerization yield (lower intensity of the band from lane 4). Obviously the second reason lies in the not complete recycling efficiency. However, it was still quite promising to show that the PCR reagents can be polymerized, de-polymerized, and re-polymerized in a closed loop. Without adding new DNA templates (Figure 2b, lane 5), there was no 2^nd^ PCR product obtained, showing that all the added DNA template and the residue of the 1^st^ PCR product were eliminated by enzymatic hydrolysis without bringing contamination to the new circle of PCR. The whole circulation of poly-oligo/mono nucleotides was also processed by using primers without PS tail protection as negative control. There was 1^st^ PCR product (lane 6), hydrolyzed PCR product (lane 7) but no 2^nd^ PCR product observed (lane 8), showing that without terminal PS-protection, primers were hydrolyzed during the nucleotide recycling step.

These results show that by using PS-modified primers it is possible to circulate poly-oligo/mono nucleotide materials in PCR kit and repeat use the PCR substrate in a closed loop. The system is still far from ideal. For example, we can mention that in the recycled PCR kit there is an addition band beside the expected band of PCR amplicons (∼100 bp, lane 4 and 5). We believe that this band is probably the result of cleaved PCR amplicons with shorter lengths (Figure S3). In support of this conclusion, we should mention that when the recycled primers were purified by Oligo extraction kit and applied for another round of PCR amplification, the extra band was absent (Figure S4).

### Recycle primers for Qpcr

We show that the nucleic acid materials in the PCR kit can be recycled by using PS-modified primers. However, for the detection of specific DNA, an additional step of gel electrophoresis is required to show the band of the amplicon. This ‘end point’ PCR is time-consuming and lacks efficiency. For molecular diagnostic, quantitative PCR (qPCR) is widely used as DNA amplification can be monitored in real-time by measuring the fluorescence signal of DNA intercalation dye. For qPCR in molecular diagnostic, primers are critical components, as their functions of recognizing and binding with target DNA decide the precision and sensitivity of the molecular diagnosing assay.^28^ Consequently, in this circular recycling work the quality of recycled primers (purity and sequence length) would directly affect the performance of qPCR. Above we have shown that 4PS-modification can protect the primers from enzymatic hydrolysis, but the recycled primers have different lengths (with 0 to 3 nucleotides loss) this in theory should lead to the use of a more complex annealing temperature profile for an optimized recycled PCR use.

To maintain the length of the recycled primers without changing too much of their annealing temperature, primers with a 2-PS tail at 3’-terminal modification were used for qPCR (amplicon length 133 nucleotides) and recycled by the above-mentioned method. The function of 2PS-modified primers binding to their DNA targets and the polymerase catalyzed chain extension was evaluated by qPCR. The qPCR performance was not affected by primers with 2PS-modification (Figure S5a). The 2PS-modified primers also showed very good nuclease hydrolysis tolerance (Figure S5b).

Agilent Oligo Pro II system was applied to monitor the hydrolysis process for the recycling of 2PS-modified qPCR primers. As a capillary electrophoresis analytical tool, Oligo Pro II is suitable for the detection of single-strand oligonucleotide (sub 60 nt) with single nt resolution. The qPCR product was incubated with a mixture of nucleases for different times (1h, 2h, and 4h) to hydrolyze the amplicon and to recycle the primers. The recycled primers were then purified by the Oligo extraction kit and analyzed by Oligo Pro II, with a mixture of fresh primers used as a control. As shown in Figure 3, the forward and reverse primers (20 nt and 23 nt, respectively) migrated for different times and thus can be separated well. The difference in the peak intensity between the forward and reverse primers is due to the differences in their sequences (i.e. base content). As shown in Figure 4a, for the recycled primers, after 1h incubation with nucleases, there were still several longer oligonucleotides observed. In the plot these are peaks 10-20 whose migration time is longer than that of the fresh primers (Figure 3) that in Figure 4a should be peaks 7 and 9. These longer oligonucleotides should be residues of qPCR amplicons. In Figure 4a one can also observe a few very short oligonucleotides residue (peaks 1-5) due to the incomplete hydrolysis. After a longer incubation time (2h), the longer oligonucleotides residues were not detected anymore (Figure 4b), showing that the qPCR residues can be fully eliminated by nuclease hydrolysis. There are four major peaks observed from the primer recycling products (peak 5-8, Figure 4b), which are recycled primers with fully preserved lengths and reproducible migration times as fresh primers (peak 6 and 8), and recycled primers with one terminal extra nucleotide hydrolysed showing shortened migrations time (peak 5 and 7). The amount of very short oligonucleotides residue was also decreased with prolonged hydrolysis time (1h incubation, peaks 1-5, Figure 4a; 2h incubation, peaks 1-4, Figure 4b; 4h incubation, peaks 1-2, Figure 4c). These results show that 2PS-modification can effectively protect the primers from nuclease hydrolysis. The recycled primers can either be fully preserved or loose 1nt. Further, the mixture of recycled primers was characterized by MALDI-TOF. Similar as the Oligo Pro II characterization, fully preserved primers, and primers with only one terminal nucleotide lost were obtained (mass spectra in Figure 5a).

**Figure 3.**
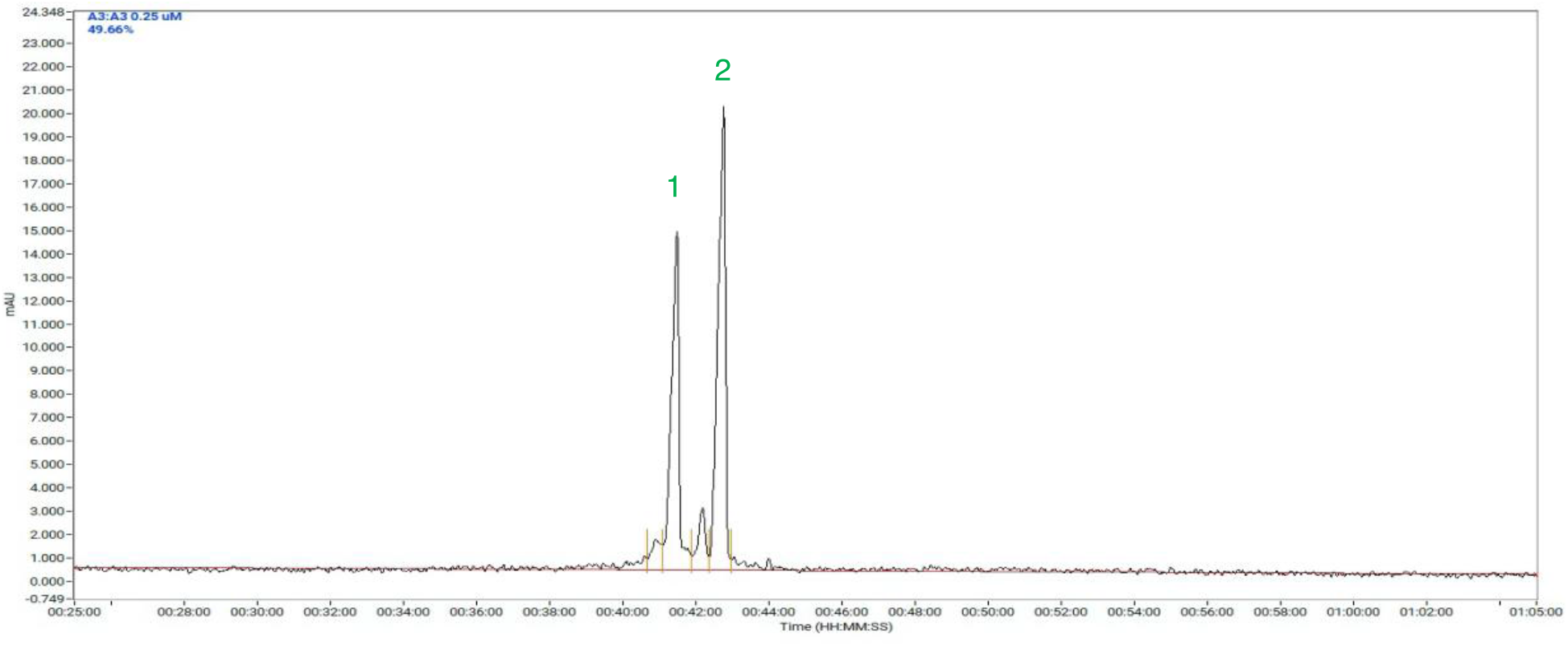
Characterization of qPCR primers by Oligo Pro II capillary electrophoresis. Retention time of fresh forward (peak 1) and reverse (peak 2) primers with 3’ terminal 2PS-modification.

**Figure 4.**
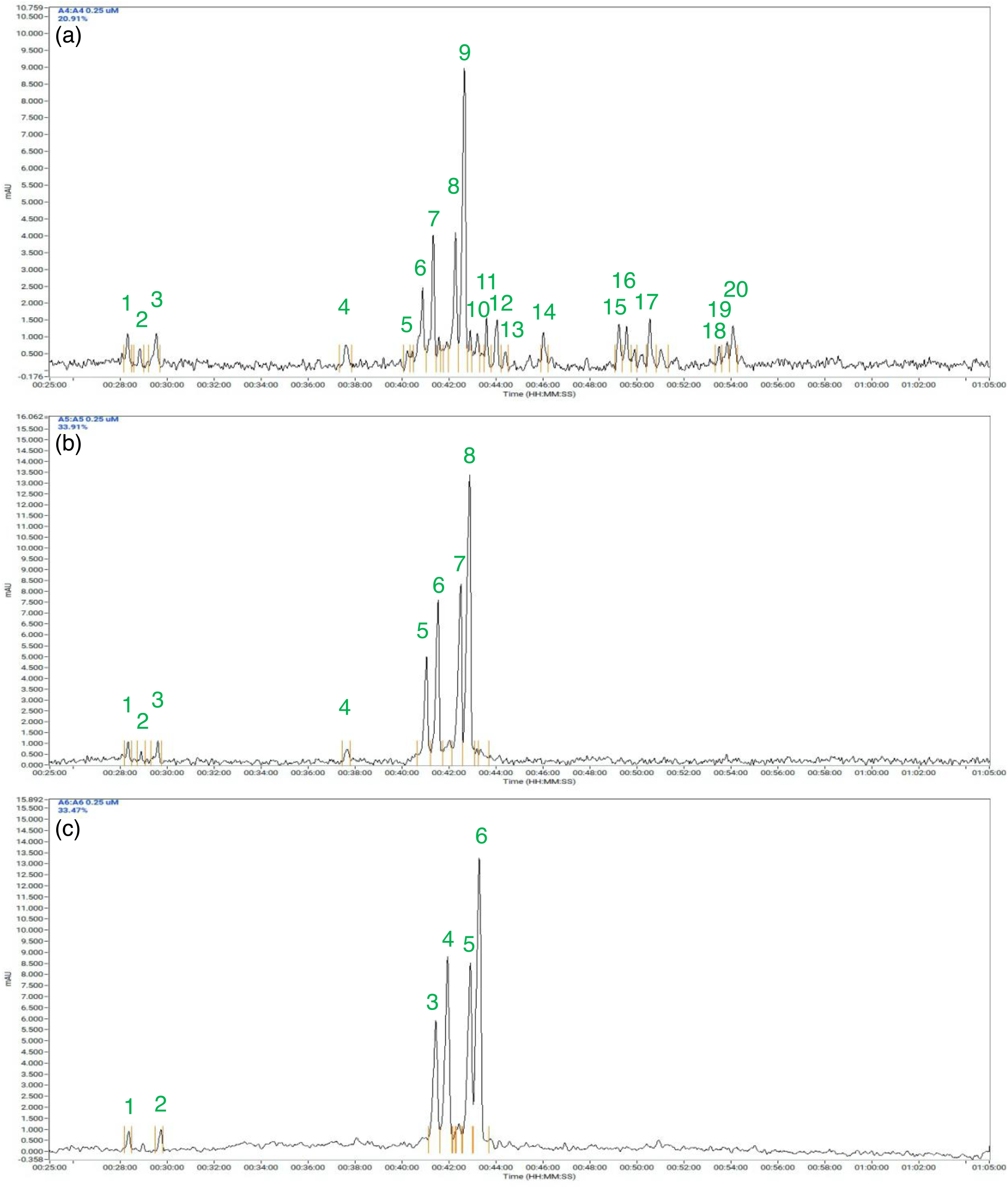
Monitoring the hydrolysis process of qPCR product as well as the recycling process of primers by Oligo Pro II capillary electrophoresis. Retention time of recycled primers from qPCR kit after (a) 1h, (b) 2h, and (c) 4h hydrolysis by mixture of nuclease, the primers were purified by Oligo extraction kit before injection into the Oligo Pro II capillary electrophoresis system.

**Figure 5.**
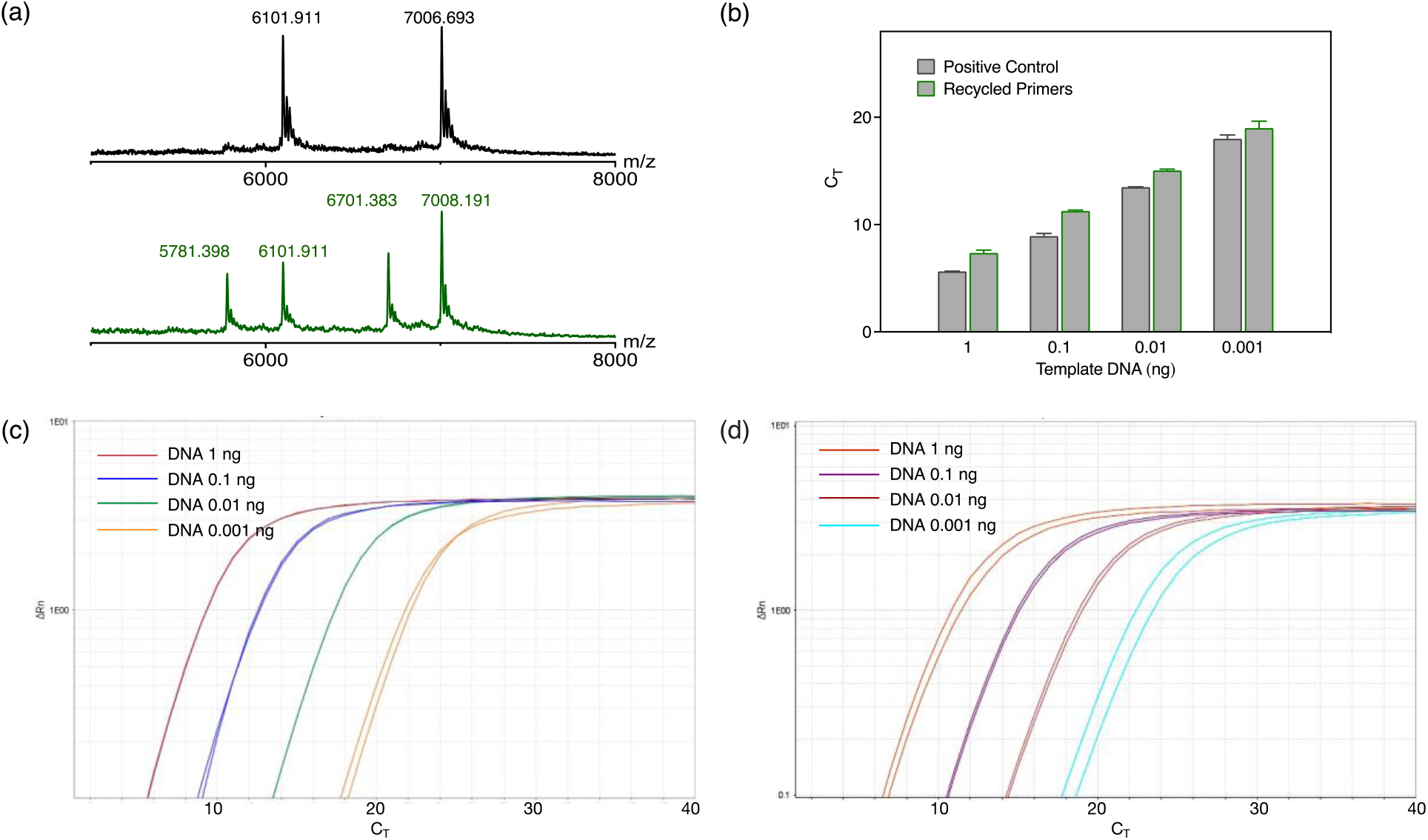
(a) MALDI-TOF characterization of fresh and recycled primers with 2 PS-tail modification. (b) CT value of qPCR amplification with fresh and recycled primers DNA template was 1, 0.1, 0.01, 0.001 ng, respectively. (c) qPCR amplification plots with fresh primers. (d) qPCR amplification plots with recycled primers.

The recycled primers were applied for qPCR amplification, with fresh primers used as positive control (Figure 5). Similar CT values were obtained of the qPCR assay prepared with fresh primers and recycled primers (Figure 5b-d), showing that the recycled primers can be circularly used as new materials for qPCR amplification and DNA testing. The recovery yield of qPCR primers is 40.93 ± 0.48 %. Due to the loss during oligo extraction, we assume the preserved primers should be more than 50 %. Even with only 2-PS modification, the qPCR primers can be effectively preserved and circularly re-used in qPCR assay without decreasing the detection sensitivity too much (less than 2 thermo-cycles). For the NTC sample (no template control), the CT value 22 is lower than the control sample (32 thermo-cycles, Figure S7b) indicating the presence of unwanted residues. Although this NTC result should not affect the detection limit of the qPCR kit being higher than CT value found for 0.001 ng of DNA template with the recycled primers (20, see Figure 5b), it would be better to purify the recycled primers on a larger scale (for example by ion-exchange chromatography) to eliminate the residue of DNA amplicon as well as any other potential contaminants. Overall, in case of industrial scale application, a higher yield of primers with better purities should be anticipated.

## CONCLUSION

In this work, we discussed the possibility to circularly recycle PCR kits. By using PS modification, the primers can be protected from the nucleotide recycling process, preserved, and extracted for circular usage. Together with the recycled monomeric nucleotides with good yield, the PCR substrate was regenerated and reused for the detection of targeted DNA. The PS-modified primers can also be recycled separately by the same method and reused as new materials for the targeted DNA detection by qPCR with relatively good sensitivity. The quality (length and purity) of recycled primers can be evaluated by Oligo pro II capillary electrophoresis with single nt resolution. The method we established here shows the possibility of using PCR reagents in a smarter way. Overall, the daily accumulated PCR waste during the current pandemic could become a valuable resource for the circularly recycling of PCR reagents. With the PS-modification it is possible to design non-hydrolysable, recyclable primers to ease the problem of PCR kit shortage. There should be also a chance to improve the recycling efficiency of primers by using ‘stereopure’ PS-modification in the future.^27^ So far the established method was tested by end point PCR and qPCR (with intercalation dye), but it might also be applicable to recycle oligo-and mononucleotides from the other DNA polymerization system like LAMP system (Loop-mediated isothermal amplification).^29^ As DNA technology has become a very well-established technique for sequencing, bioengineering, and molecular diagnoses, this DNA recycling methodology could also be easily adapted for the circularly recycle of modified nucleotides (e.g., fluorescence dye-labeled dNTPs for sequencing^30^), as well as oligonucleotide products (e.g., DNA microarray^31^).

## METHOD

### Materials

#### Chemicals and consumables

Ethanol, Isoproponal, and 2-mercaptoethanol (for oligo-extraction) were purchased from Sigma-Aldrich. 2’-Deoxyguanosine 5’-monophosphate sodium salt hydrate (dGMP), Thymidine 5′-monophosphate disodium salt hydrate (dTMP), 2′-Deoxycytidine 5′-monophosphate sodium salt (dCMP), 2′-Deoxyadenosine 5′-monophosphate (dAMP), Acetylephosphate Lituium Potassium salt (AceP), Sodium chloride (DNase, RNase, and protease free), Sodium hydroxide solution (5.0 M), Acetic acid, Tetrabutylammonium dihydrogenphosphate solution (1.0 M in water), Deoxynucleotide mix reagent (dNTPs, 10 mM for each), Adenosine 5’-triphospate disodium (ATP, 100 mM) were purchased from Sigma-Aldrich. Nuclease-Free Water (10 × 50 ml) was purchased from QIAGEN. *Self-made phosphorylation buffer:* 55mM HEPES, Magnesium acetate 15 mM, pH 7.5. HEPES buffer (1 M) was purchased from Thermo Fisher Scientific. *Chemicals for gel electrophoresis*: 400 μL-SYBR Safe DNA Gel Stain, DNA Loading Dye and SDS Solution (6x), TAE Buffer (Tris-acetate-EDTA) (50x), TrackIt 100 bp DNA Ladder, Ultra Low Range DNA Ladder were purchased from Thermo Fisher Scientific. Agarose was purchased from Bio-Rad Laboratories. NuPAGE™ 4 to 12%, Bis-Tris, 1.0 mm, Mini Protein Gel, 12-well was purchased from Invitrogen. *PCR and qPCR reagents:* DreamTaq Green PCR Master Mix (for first round of PCR), DreamTaq polymerase (for second round of PCR), and PowerTrack SYBR Green Master Mix (for qPCR) were purchased from Thermo Fisher Scientific. Amicon Ultra-0.5 mL Centrifugal Filters (3KD cutoff) was purchased from Merck. Oligo Clean-Up and Concentration Kit was purchased from Norgen Biotek. *Enzymes*: Restriction enzymes HpyF3I (DdeI), BshNI (BanI), PsuI (BstYI), Kpn2I (BspEI) were purchased from Thermo Fisher Scientific. Hydrolysis enzymes Exonuclease III (200 U/μL) and Exonuclease I (20 U/μL) were purchased from Thermo Fisher Scientific. T4 NMP (Nucleotide Monophosphate) Kinase was purchased from Jena Bioscience. E. coli S30 Extract System (with luciferase plasmid DNA template) was purchased from Promega Corporation. *DNA template and primers for PCR*: Plasmid luciferase DNA (pBEST luc, 4864 bp) was from the E. coli S30 extraction kit. Primers were designed from published sequence^24^ without 5′ NdeI and 3′ BamHI restriction sites sequence. The amplification length is 1653 base pair (16 to 1668) of the luc sequence. Primers (oligonucleotides) with or without 5’-terminal Fam modification, and with or without 3’-phosphorothiolate (PS) modification, were purchased from Biomers Gmbh, Germany. The 3-terminal PS tail was labeled by ‘*’ in the flowing sequences. (Forward primer (Fam) 5’-atg gaa gac gcc aaa aac at*a *a*a*g-3’, Reverse primer (Fam) 5’-tta caa ttt gga ctt tcc gc*c *c*t*t-3’). *Primers for quantitative PCR (qPCR)*: DNA template for qPCR is the same as PCR (luc template). Primers were designed by IDT PrimerQuest™ Tool with forward primer starts from 673 of luciferase sequence, reverse primer start from 805 of luciferase sequence, and length of amplicon 133 base pair (the same template and primer sequences as qPCR in DNA NaCRe system^5^). Primers were ordered from Biomers Gmbh. The 3-terminal PS tail was labeled by ‘*’ in the flowing sequences (Forward primer (Fam) 5’-cgc atg cca gag atc cta *t*t -3’, Reverse primer (Fam) 5’-aga cga ctc gaa atc cac ata *t*c -3’).

### Instruments

Eppendorf ThermoMixer (RTM F1.5, 220 - 240 V/50 - 60 Hz) was purchased from Eppendorf. Horizontal gel electrophoresis system was purchased from Bio-Rad. Gel images was taken from GelDoc Go system, Bio-Rad. PCR was performed by Proflex 3×32-well PCR thermal cycler system (Thermo Fisher Scientific). qPCR was performed by QuantStudio 7 qPCR instrument (Applied Biosystems). HPLC was performed by Infinite 1260 HPLC with C 18 column, Agilent.

#### Thermocycle condition for PCR

initial denature (95°C, 2 min) for 1 cycle, amplification (denature at 95°C for 30s, annealing at 55 °C for 30 s, extension at 72 °C for 1 min) for 35 cycles, final extension (72 °C for 10 min) for 1 cycle and cooling to 4°C. The PCR amplification products were used for recycling of dNTPs, re-generation of PCR kit, and recycling of primers. Unless specified, all the PCR were processed under the same thermocycle condition.

#### Gel electrophoresis

The agarose gel was run in 1 × TAE buffer at 120 V for 40 min. Afterwards, the gel was stained by 1x Sybr safe solution for 40 min under slow shaking. Following the gel image was taken by GelDoc Go under Sybr safe channel for 1 s exposure time (Figure 1b). Unless specified, all agarose gel experiments were processed under the same condition.

#### Thermocycle condition for qPCR

initial denature (95°C, 2 min) for 1 cycle, amplification (denature at 95°C for 30s, annealing at 55 °C for 30 s, extension at 72 °C for 1 min) for 35 cycles, final extension (72 °C for 10 min) for 1 cycle and cooling to 4°C. The PCR amplification products were used for recycling of dNTPs, re-generation of PCR kit, and recycling of primers. Unless specified, all the PCR were processed under the same thermocycle condition.

#### MALDI-TOF

All the MALDI-TOF spectra were collected using a Bruker AutoFlex Speed instrument (Bremen, Germany).

#### Oligo-Pro II capillary electrophoresis

Agilent Oligo Pro II system with DN-415 OLIGEL ssDNA Gel using the corresponding methods.

## Experiments

### 1. Recycling of dNTPs from PCR products/waste

#### 1.1. Nuclease resistance evaluation of 4PS-modified primers

##### Nuclease-resistance test of PS modified primers

The nuclease-resistance capacity of PS modified primers was tested by mixing Fam-FP-PS, Fam-RP-PS, Fam-FP, Fam-RP (0.5 μL, 100 μM) with Exonuclease III (5 μL, 500 Unit) and Exonuclease I (5 μL, 50 Unit), 10 μL of 10x Exonuclease III buffer, 79.5 μL of nuclease-free water (in total 100 μL), and the nuclease hydrolysis mixtures were incubated at 37°C 350 RPM for 2, 6 and 24h. Further the hydrolysis reaction mixtures (5 μL for each) were loaded to an 2% agarose gel. As shows in the gel (Figure S1b), the FAM dye labeled, forward and reverse primer with PS protection (Fam-FP-PS and Fam-RP-PS) can tolerate enzymatic hydrolysis for 24h (lane 1-3 (Fam-FP-PS) and lane 7-9(Fam-RP-PS)). In contrast, the forward and reverse primer without PS tail (Fam-FP and Fam-RP) were hydrolyzed with very less residues after 2h (lane 6 (Fam-FP) and lane 9(Fam-RP)). This shows the primers with 4PS tail modification can resist nuclease catalyzed hydrolysis.

#### 1.2. PCR amplification and recycling of dNTPs

##### PCR reaction mixture was prepared as following

DreamTaq master mix (2x, 50 μL), forward primer (0.5 μL, 100 μM), reverse primer (0.5 μL, 100 μM), Luc DNA template (0.5 μL, 100 ng/μL), nuclease-free water (48.5 μL) in total 100 μL. The PCR products (triplicate, primers with or without 4PS-modification) were extracted by PCR extraction kit, and the concentration of PCR products was quantified by Nanodorp.

##### Cleavage and Hydrolysis

The PCR product was firstly heated to 100°C and incubated for 15 min to deactivate polymerase. Further 50 μL of PCR product was mixed with 3.25 μL Dedl (32.5 Unit), 4 μL Exo III, 4 μL Exo I, and incubated at 37°C, 350 pm overnight. At the cleavage site of Dedl (C^TNAG, 629 and 1043 of luc sequence), 5’-terminal overhangs with length of 4 nucleotides were generated, which is suitable for the binding of Exo III to the cleaved PCR product and initiation the hydrolysis. Afterwards the hydrolysis mixture was heated to 80° and incubated for 15 min to in activate the restriction and hydrolysis enzymes. Due to the increased volume by adding the cleavage and hydrolysis enzymes there was a 1.225x dilution for the reaction mixture.

##### Phosphorylation

20 μL of PCR product hydrolysis mixture was mixed with 1.2 μL E. coli S30 Extract (20x dilution), 1 μL T4 dNMP Kinase (50x dilution, 2 Unit), 1 μL ATP (3 mM), and 0.77 μL Acetylphosphate Lithium potassium (AceP, 50 mM), in total 24 μL. The phosphorylation reaction mixture with final volume 24 μL, estimated dNMPs (in average about 0.15 mM for each), ATP (0.125 mM), AceP (1.6 mM, 1.5 equivalent) was incubated in thermomixer at 400 RPM, 37°C for 4 hours. Afterwards all hydrolysis enzymes and phosphorylation enzymes, and non-hydrolyzed DNA was removed by ultrafiltration (Amicon, 3 KD cutoff, 5000 RPM for 10 min in 4°C). Due to the increased volume by adding the phosphorylation reagents there was a 1.2x dilution for the reaction mixture.

#### 1.3. Quantification of recycled dNTPs by HPLC

The concentration of recycled dNTPs was quantified by HPLC with a C-18 reverse-phase column. Mobile phase Buffer A: 5 mM t-butyl ammonium phosphate, 10 mM KH2PO4, and 0.25% methanol adjusted to pH 6.9. Buffer B: 5 mM t-butyl ammonium phosphate, 50 mM KH2PO4, and 30% methanol (pH 7.0). From 0 to 15 mins the gradients changed from 40%/60% to 20%/80% of buffer A/B and run under the same gradient condition until 20 min, and changed back to the starting condition of 40%/60%, with flow rate at 0.5 ml/min.

##### HPLC quantification

Further the filtrated reaction mixture dNTPs_post PCR, dNTPs_post hydrolysis, dNTPs_post phosphorylation was diluted (50x) and injected to HPLC (50 μL) for the quantification of dNTPs. The mixture of dNMPs (2.5 μM for each), and dNTPs from original PCR kit (100x dilution, 4 μM for each) was injected to HPLC (50 μL) as positive control. Retention times of the above five samples see Figure S2a. Calibration curve achieved by a series dilution of dNTPs and ATP (2.5-40 μM, Figure S2b). The final concentration of dNTPs was multiplied with the dilution factors during the PCR kit regeneration process (1.225x post hydrolysis, and 1.2x for phosphorylation). After PCR amplification, about 70% of dNTPs was consumed (Figure S2c and d). In addition to that, there were new peaks generated (Figure S2a_dNTPs_post PCR), which were attributed to dNTPs hydrolysis products (dNMPs and dNDPs) probably induced by heating in PCR condition. After enzymatical hydrolysis the amount of dNMPs was highly increased (Figure S2a-post hydrolysis, retention time of extra fractions between 3-14 mins). There is a slightly decrease of dNTPs residue, that might be caused by may enzymatical hydrolysis. After Phosphorylation, the dNMPs peaks were almost disappeared, and the dNTPs peaks were re-generated, showing the very good phosphorylation efficiency (Figure S2a_post phosphorylation). The recycling efficiency of dNTPs is 74.49 ± 2.09 % with the final concentration of all dNTPs is 101.35 ± 2.84 μM (Figure S2c and d).

### 2. Recycling of PCR kit

#### 2.1. PCR amplification and re-generation of PCR substrate

##### PCR amplification of luc DNA

PCR amplification was the same as above mentioned ratio of DNA template, and primers. Although with relatively good monomers recycling efficiency, we cannot exclude all the PCR residues has been removed without bring any contaminate for the next round of PCR. To improve the hydrolysis efficiency, we further separated the hydrolysis of PCR product into two steps with the restriction enzymes only added at the second step.

##### Cleavage and Hydrolysis

The PCR product was firstly heated to 100°C and incubated for 15 min to deactivate polymerase. Further 50 μL of PCR product was mixed with 1.5 μL Exo III, 1.5 μL Exo I, and incubated at 37°C, 350 pm overnight. Afterwards the hydrolysis mixture was heated to 80°C and incubated for 15 min to in activate the hydrolysis enzymes. Next, 1 μL DedI (10 Unit), 1.5 μL Exo III, 1.5 μL Exo I, were added to the hydrolysis mixture and incubated at 37°C, 350 pm for 8h. Afterwards the cleavage and hydrolysis mixture was heated to 80°C and incubated for 15 min to in activate the restriction and hydrolysis enzymes. Due to the increased volume by the two steps of hydrolysis and cleavage there was a 1.12x (50 to 56 μL) dilution for the reaction mixture.

##### Phosphorylation

The phosphorylation step was processed similar as above mentioned method. As the phosphorylation mixture would add to the total volume of the reaction mixture and decrease the concentration of all the reagents in the PCR substrate, we would like to minimize the added phosphorylation mixture. To do that, we first prepared a phosphorylation reagents mixture (10 μL) with T4 60x dilution, S30 20x dilution, ATP 2.46 mM, AceP 31.5 mM. 30 μL of the PCR hydrolysis mixture was mixed with 2 μL of the prepared phosphorylation reagents, and the phosphorylation mixture was incubated in thermomixer at 400 RPM, 37°C for 4 hours. Afterwards the reaction mixture was directly applied for PCR amplification. Due to the increased volume by adding the phosphorylation reagents there was a 1.066x dilution (30 to 32 μL) for the reaction mixture. Calculated from the recycling efficiency of monomers, as well as the dilution factor for each step of PCR kit re-generation, the final concentration of dNTPs in the re-generated PCR kit is as following: Conc.dNTPs * Recycling % / dilution factors = 200 μM * 74.49% /1.12/1.066/1.05 = 118.84 μM

#### 2.2. Recycling of re-generated PCR substrate

##### PCR reaction mixture was prepared as following

20 μL of re-generated PCR substrate, 0.5 μL of luc DNA template (20 ng/μL), 0.5 μL of DreamTaq polymerase (diluted to 1U/ μL), in total 21 μL. *No DNA template control was prepared as following:* 20 μL of re-generated PCR substrate, 0.5 μL of nuclease-free water, 0.5 μL of DreamTaq polymerase (diluted to 1 U/ μL), in total 21 μL. Due to the increased volume by adding the DNA template and the polymerase there was a 1.05x dilution (21 to 20 μL) for the reaction mixture. The whole process for PCR amplification, re-generation, second round of PCR was processed by primers without PS modification as a negative control.

##### Gel electrophoresis

after the PCR thermocycles, the first and second round of PCR product, as well as PCR-hydrolysis mixture, were loaded to an 2% agarose gel. Samples were all mixed with 2 μL of 6x DNA loading dye. lane 1, TrackIt 100 bp ladder, 2 μL; lane 2, first round of PCR product 8.4 μL + 1.6 μL water; lane 3, PCR-hydrolysis mixture, 9.375 +0.625 μL water; lane 4, second round of PCR product with DNA template added, 10 μL; lane 5, second round of PCR product without DNA template added, 10 μL. Lane 6 – lane 8 first and second round of PCR product (primers without PS modification), lane 6, first round of PCR product 8.4 μL + 1.6 μL water; lane 7, PCR-hydrolysis mixture 9.375 μL + 0.625 μL water; lane 8, second round of PCR product with DNA template added, 10 μL.

### 3. Recycling of primers

#### PCR amplification of luc DNA

PCR amplification (duplicate) was the same as above mentioned ratio of DNA template, and primers. The concentration of primers was quantified by Nanodrop (ssDNA method, 1433.05 ng/μL at 100 μM), and added primers was 716.5 ng (0.5 μL, 100 μM).

#### Cleavage and Hydrolysis

The PCR product (100 μL) was firstly heated to 100°C and incubated for 15 min to deactivate polymerase. Further the PCR product was mixed with BshNI (BanI) (2.5 μL, 25 Unit) At the cleavage site of BshNI (BanI) (G^GYRCC, 48 of luc sequence), 5’-terminal overhangs with length of 4 nucleotides were generated. Another restriction enzyme PsuI (BstYI) (10 μL, 100 Unit) with multiple cleavage sites (R^GATCY, 613, 684, 1137, 1625 of luc sequence) was also added to cleave the PCR product. Also, the cleaved fragment with forward primer was shortened to 33 nt (16-48), and the cleaved fragment with reverse primer was shortened to 43 nt (1625 to 1668) for better hydrolysis and recycle efficiency. The cleavage mixture was incubated at 37°C, 350 rpm for 5 h. Afterwards the hydrolysis mixture was heated to 80 °C and incubated for 20 min to inactivate the cleavage enzymes. Next, 15 μL of Exo III, 15 μL of Exo I were added to the hydrolysis mixture and incubated at 37°C, 350 rpm for 8 h. Afterwards the hydrolysis mixture was heated to 80°C and incubated for 15 min to inactivate the hydrolysis enzymes. Further, the PCR-cleavage-hydrolysis mixture was concentrated by ultrafiltration (3K cutoff, 8000 rpm, 45 min, 4 °C) to 50 μL. Mixture of primers was extracted by ‘Oligo Clean-Up and Concentration Kit’. The concentration of extracted primers was quantified by Nanodrop (ssDNA method) with final amount of 436 ± 7.2 ng (40 μL, 10.90 ± 0.18 ng/ μL). *Recycling yield (mass ratio):* 436 ± 7.2 ng / 716.5 ng = 60.85 ± 0.01 %.

#### MALDI-TOF

The molecular mass of fresh primers and recycled primers were characterized by MALDI-TOF. Matrix was prepared as following: 90:10 mixture of (1) 50 mg/ml 3-Hydroxy picolinic acid in 1:1 water/acetonitrile, and (2) 100 mg/ml diammonium hydrogen citrate in water. ^19^ All the MALDI-TOF spectra were collected using a Bruker AutoFlex Speed instrument (Bremen, Germany). Forward primer (2.5 μM in water), reverse primer (2.5 μM in water), and recycled primers (10.90 ± 0.18 ng/μL in water) were mixed with an equal volume of matrix solution. For each sample, 1 μL aliquot of such solution mixture was deposited and dried onto a stainless ground steel target plate. Measurements were performed in positive ionization mode and operated in the linear mode in the 1K - 14K m/z mass range. The laser intensity was kept at around 80 % for all measurements. Typically, around 1000 shots were accumulated for each spectrum. Mass spectra were processed with Flex Analysis (Bruker) software.

#### Recycling yield (molar ratio)

Due to the shortened length of primers, the molecular weight of recycled primers was decreased from 7.5 KD and 7.3 KD to 6.9 KD (in average). Therefore, the recycling yield was calculated by molar ratio is 64.38 ± 0.01%. The so achieved concentration of recycled primers was 0.82 μM, calculated from the molar ratio.

#### PCR reaction mixture was prepared as following

(1) Positive control of PCR by fresh primes: DreamTaq master mix (2x, 10 μL), mixture of forward and reverse primers (8 μL, 1.25 μM), Luc DNA template (1 μL, 10 ng/μL), nuclease-free water (1 μL) in total 20 μL. (2) PCR with recycled primers: DreamTaq master mix (2x, 10 μL), recycled primer mixture (8 μL, ∼ 0.82 μM) Luc DNA template (1 μL, 10 ng/μL), nuclease-free water (1 μL) in total 20 μL. The thermocycler condition is the same as above settings. After the PCR amplification, PCR products from fresh and recycled primers (10 μL for each) were loaded to a 2% agarose gel.

### 4. Recycle primers for qPCR

#### 4.1 Nuclease resistance evaluation of 4PS-modified primers

##### Nuclease-resistance test of PS modified primers

The nuclease-resistance capacity of PS modified primers was tested by mixing (0.5 μL, 100 μM) of Fam-FP-PS, Fam-RP-PS, Fam-FP, Fam-RP with Exonuclease III (5 μL, 500 Unit) and Exonuclease I (5 μL, 50 Unit), 10 μL of 10x Exonuclease III buffer, 79.5 μL of nuclease-free water (in total 100 μL), and incubation overnight at 37°C, 350 RPM. Further the hydrolysis reaction mixtures (5 μL for each) were loaded to an 2% agarose gel. As shows in the gel (Figure S1), the FAM dye labeled, forward and reverse primer with PS protection (Fam-FP-PS and Fam-RP-PS) can tolerate enzymatic hydrolysis (lane 2 and lane 4). There is no obvious difference of the bands in comparison to the control samples of nuclease free primers (lane 1 and lane 3). In contrast, the forward and reverse primer without PS tail (Fam-FP and Fam-RP) were hydrolyzed with very less residues (lane 6 and lane 8) in comparison to the control samples of nuclease free primers (lane 5 and lane 8). This shows the PS tail modification of primers could resist nuclease catalyzed hydrolysis.

As qPCR experiments are normally processed in very small scale (10 or 20 μL for each sample). Next, we tried to recover primers from PCR product/waste (100 μL scale for qPCR). *PCR reaction mixture was prepared as following:* PowerTrack SYBR Green Master Mix (2x, 50 μL), mixture of forward and reverse primers (1.6 μL, 25 μM) with final concentration 400 nM, Luc DNA template (1 μL, 1 ng/μL), nuclease-free water (47.4 μL) in total 100 μL (duplicate, with 4 × 100 μL in each group). The concentration of primers was quantified by Nanodrop (ssDNA method, 361.12 ng/μL at 25 μM), and added primers was 577.79 ng /100 μL of PCR mixture (1.6 μL).

##### Thermocycle condition for PCR

initial denature (95°C, 2 min) for 1 cycle, amplification (denature at 95°C for 5 s, annealing and amplification at 60 °C for 15 s) for 35 cycles, final extension (72 °C for 10 min) for 1 cycle and cooling to 4°C.

##### Cleavage and Hydrolysis

The PCR product was firstly heated to 100°C and incubated for 15 min to deactivate polymerase. Further 100 μL of PCR product was mixed with Kpn2I (BspEI) (1.5 μL, 15 Unit) At the cleavage site of Kpn2I (BspEI) (T^CCGGA, 711 of luc sequence), 5’-terminal overhangs with length of 4 nucleotides were generated. The amplified DNA was cleaved into 2 shorted fragments (Figure S6a, length of 39 and 94 nt, respectively). The DNA cleavage mixture was incubated at 55°C, 350 rpm overnight. Afterwards the reaction mixture was heated to 80 °C and incubated for 20 min to inactivate the cleavage enzymes. Next, 5 μL of Exo III, 5 μL of Exo I were added to the hydrolysis mixture and incubated at 37°C, 350 rpm for 1.5 h. Afterwards the hydrolysis mixture was heated to 80°C and incubated for 15 min to inactivate the hydrolysis enzymes. Further, the PCR-cleavage-hydrolysis mixture (all the 4 × 100 μL) was concentrated by ultrafiltration (3K cutoff, 8000 rpm, 45 min, 4 °C) to 50 μL. Mixture of primers recycled from the Quadruplicates was extracted by ‘Oligo Clean-Up and Concentration Kit’. The concentration of extracted primers was quantified by Nanodrop (ssDNA method) with final amount of 873.2 ± 10.4 ng (40 μL, 21.83 ± 0.26 ng / μL, duplicate). The molecular mass of fresh primers and recycled primers were characterized by MALDI-TOF by the same method as above. *Recycling yield (molar ratio):* Due to the shortened length of primers, the molecular weight of recycled primers was decreased from 6.1 KD and 7 KD to 6.4 KD (in average, multiplied by 0.976). Therefore, the recycling yield was calculated by molar ratio is: 873.2 ± 10.4 ng / (577.79 *4) / (150/160) / 0.976 = 40.93 ± 0.48 %. Due to the lost during oligo extraction (up to 90% recovery rate with at least 10% lost), the preserved primers should be more than 50%. The so achieved recycled primers was further diluted to a stock solution (1.33 μM).

##### Monitor the primers recycling by Oligo-Pro II capillary electrophoresis

###### Sample Preparation and Methods

All samples were diluted to 0.25 uM by adding 5.0 uL of 1.0 uM stock was added to 15.0 uL nuclease-free water with mineral oil overlay. All samples were analyzed on the Agilent Oligo Pro II system with DN-415 OLIGEL ssDNA Gel using the corresponding methods. *Sample injection:* Mixture of fresh primers: 7 kV for 5 sec, sample separation: 12 kV for 90 min. Sample injection for mixture of recycled primers: 10 kV for 15 sec, sample separation: 12 kV for 90 min. *Data Analysis:* The Agilent Oligo Pro II data analysis software with the following integration parameters were used for automated data processing. The parameters were adjusted as needed to optimize the integration of individual sample in each run. The percentage purity of all the integrated peaks is 87.94% (fresh primers), 48.14% (recycled primers, 1h hydrolysis), 85.07% (recycled primers, 2h hydrolysis), 89.29% (recycled primers, 4h hydrolysis).

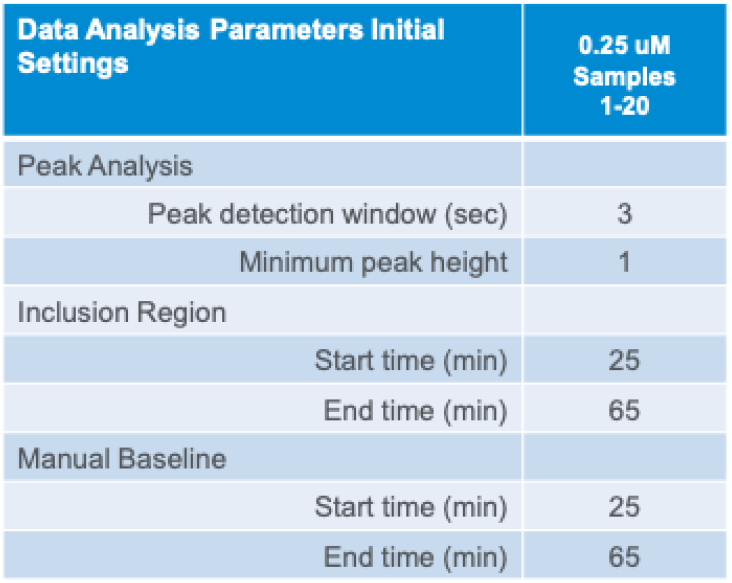

##### Duplicated qPCR mixture was prepared as following

(1) fresh primers for qPCR: PowerTrack SYBR Green Master Mix (2x, 5 μL), fresh mixture of primers (2.5 μM, 1.6 μL), luc DNA template 1 μL of series dilution with 1, 0.1, 0.01, 0.001 ng/μL and 1 μL of nuclease free water as NTC (no template control sample), 3.4 μL of nuclease-free water were mixed with in total volume 10 μL. In the final qPCR mixture, the concentration of primers is 400 nM. (2) Recycled primers for qPCR: PowerTrack SYBR Green Master Mix (2x, 5 μL), recycled primes, (1.33 μM, 3 μL), Luc DNA template 1 μL of series dilution with 1, 0.1, 0.01, 0.001 ng/μL and 1 μL of nuclease free water as NTC,1 μL of nuclease-free water were mixed with in total volume 10 μL. In the final qPCR mixture, the concentration of primers is 400 nM.

##### Thermocycle condition for qPCR

the qPCR amplification was performed by QuantStudio 7 qPCR system: initial denature (95°C, 2 min) for 1 cycle, amplification (denature at 95°C for 15 s, annealing and amplification at 60 °C for 30 s) for 40 cycles.

## Scheme and Figures

**Scheme 1.**
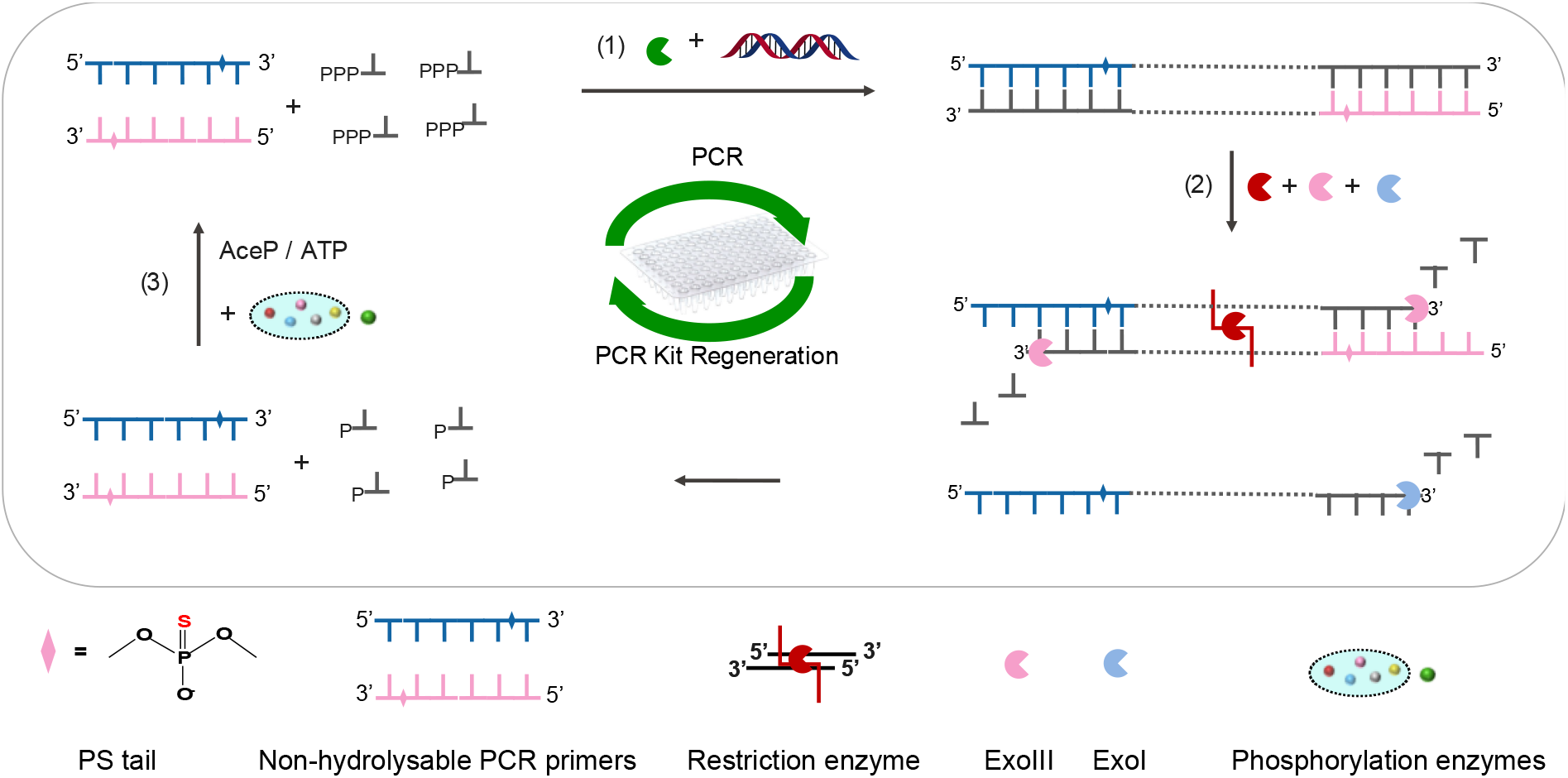
Three steps regeneration of PCR kit by using nuclease tolerant oligonucleotides with terminal phosphorothioate (PS) backbone modification as primers for PCR. Step (1) PCR amplification by using Primer-PS. Step (2) cleavage and hydrolysis of PCR product. The primers with PS-modification can tolerate enzymatic hydrolysis in this step. Step (3) one-pot phosphorylation to convert dNMPs into dNTPs.

## ASSOCIATED CONTENT

### Supporting Information

The following files are available free of charge. brief description (file type, i.e., PDF)

## AUTHOR INFORMATION

### Author Contributions

W. Liu initiated the concept of recyclable primers and PCR NaCRe system, performed the experiments, and drafted the manuscript. F. Stellacci initiated the concept of DNA NaCRe system, acquired funding, supervised the work, and revised the manuscript.

## Notes

Any additional relevant notes should be placed here.

## ACKNOWLEDGMENT

We acknowledge Agilent Technologies, Inc., and Mr. Chad Reinke (Validation scientist, Agilent) for the demo testing of fresh and recycled primers by Oligo Pro II capillary electrophoresis instrument. We thank the EPFL_ISIC group for providing the MALDI-TOF instrument and Dr. Daniel Ortiz for the training of the instrument. We thank EPFL_Core Gene Facility for providing qPCR instrument. W. Liu thanks Mr. Vincenzo Scamarcio (EPFL_SuNMiL group) for the help with the DNA hydrolysis and oligo extraction experiments.

## Supporting Figures

**Figure S1.**
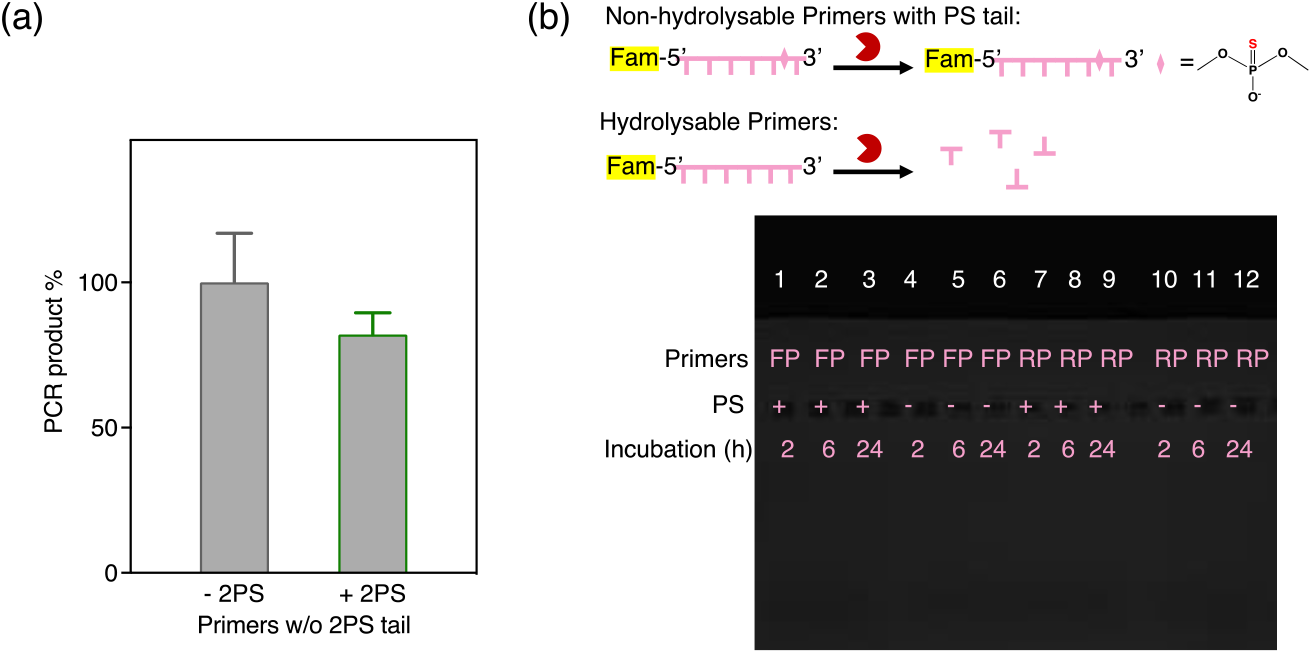
(a) Yield of PCR products by using primers with or without 4PS-modification. (b) Nucease tolerance study of primers with or without 4PS-modification. Lane 1-3, forward primer with 4PS-modification incubated with mixture of nucelases for 2, 6 and 24h. Lane 4-6, forward primer without 4PS-modification incubated with mixture of nucelases for 2, 6 and 24h. Lane 7-9, reverse primer with 4PS-modification incubated with mixture of nucelases for 2, 6 and 24h. Lane 10-12, reverse primer without 4PS-modification incubated with mixture of nucelases for 2, 6 and 24h.

**Figure S2.**
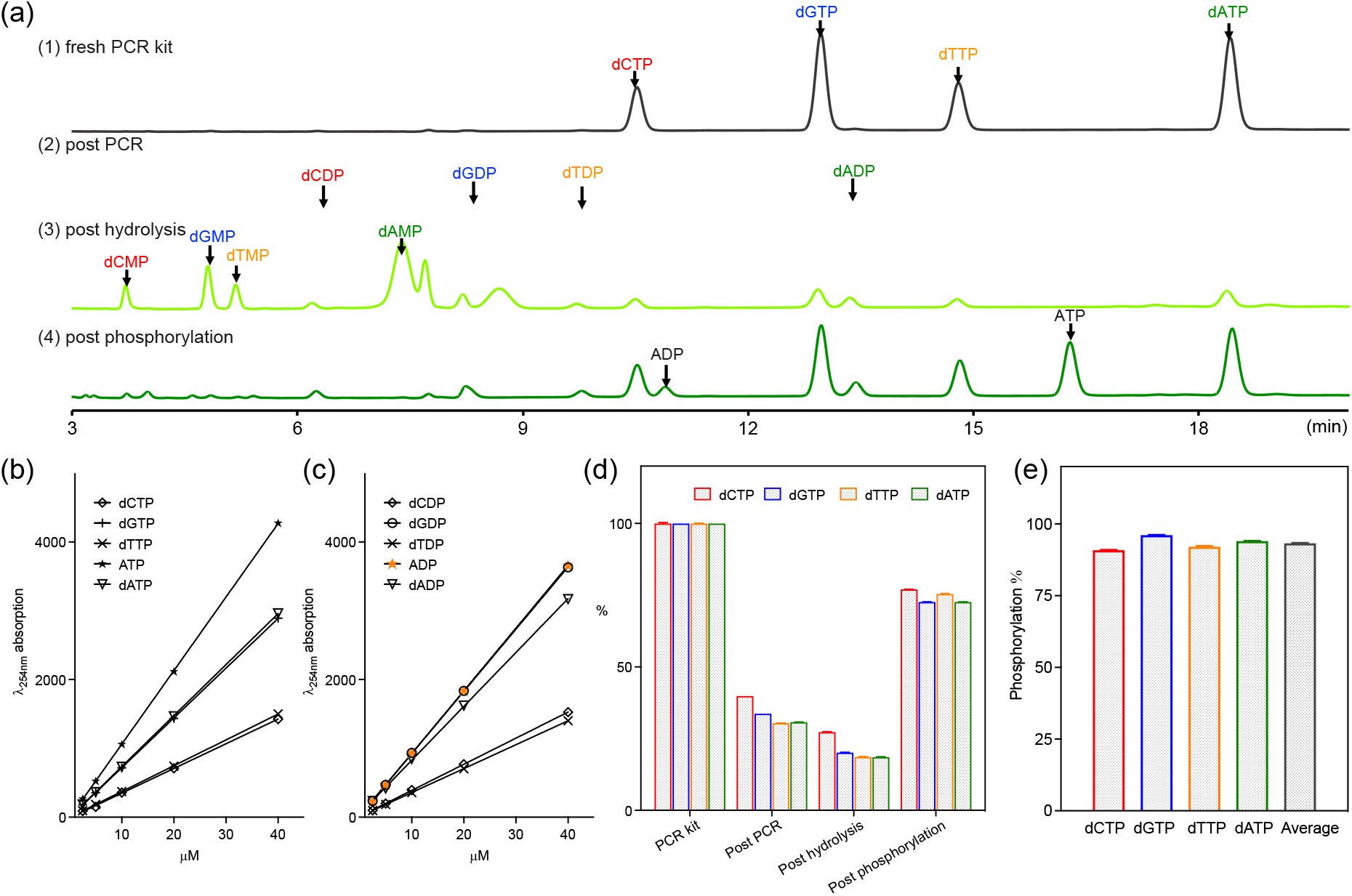
(a) HPLC retention time of monomeric nucleotides for each step of PCR kit regeneration (fresh PCR kits with dNTPs standards (line 1), dNTPs residues after PCR (line 2, post PCR), PCR product hydrolysis mixture (line 3, post hydrolysis), and regenerated PCR substrate (line 4, post phosphorylation). (b) Calibration curve of dNTPs and ATP with concentration 2.5, 5, 10, 20, 40 μM for each. (c) Calibration curve of dNDPs and ADP with concentration 2.5, 5, 10, 20, 40 μM for each. (d) Residue of dNTPs at each step of PCR kit regernation. After PCR, dNTPs were consumed and the amount of dNTPs was decreased to dC 39.86 ± 0.01%, dG 33.69 ± 0.01%, dT 30.36 ± 0.06%, dA 30.77 ± 0.03%, in average 33.67 ± 4.06%. After enzymetic hydrolysis dNTPs were partialy hydrolyzed, and the amount of dNTPs residues was decreased to dC 27.29 ± 0.11%, dG 20.15 ± 0.04%, dT 18.62 ± 0.10%, dA 18.57 ± 0.10%, in average 21.16 ± 3.84%. After phosphorylation, the dNMPs were converted to dNTPs, and the amount of dNTPs was increased to dC 77.04 ± 0.05, dG 72.64 ± 0.02%, dT 75.42 ± 0.19%, dA 72.65 ± 0.04%, in average 74.44% ± 2.01%. (e) Phosphorylation effieicency of dC 90.85 ± 0.06%, dG 96.11 ± 0.03%, dT 92.09 ± 0.23%, dA 94.00 ± 0.05%, in average 93.26 ± 2.12%.

**Figure S3.**
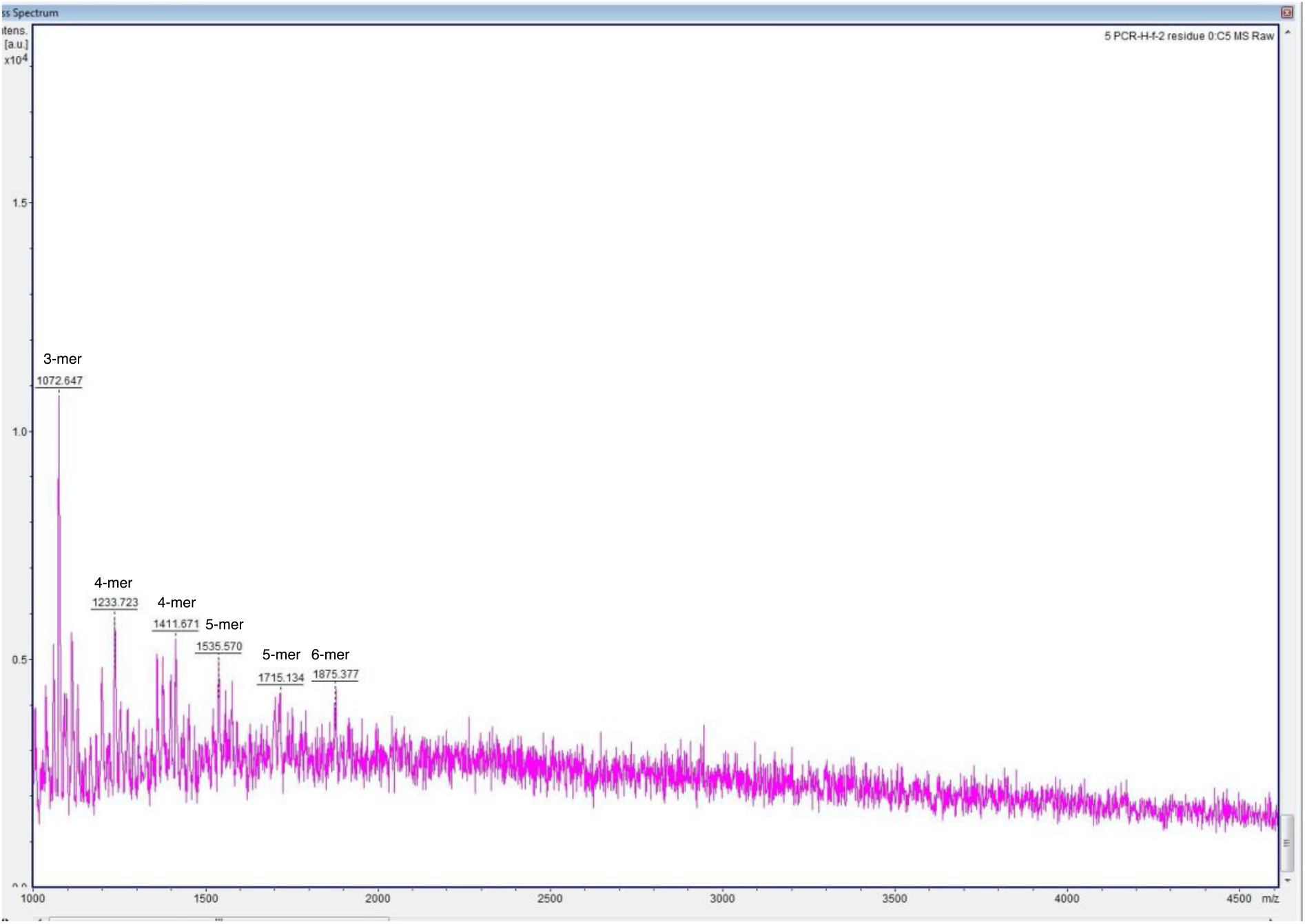
Mass spectrum of oligonucleotite residues with length of 3-6mer in the regenerated PCR kit.

**Figure S4.**
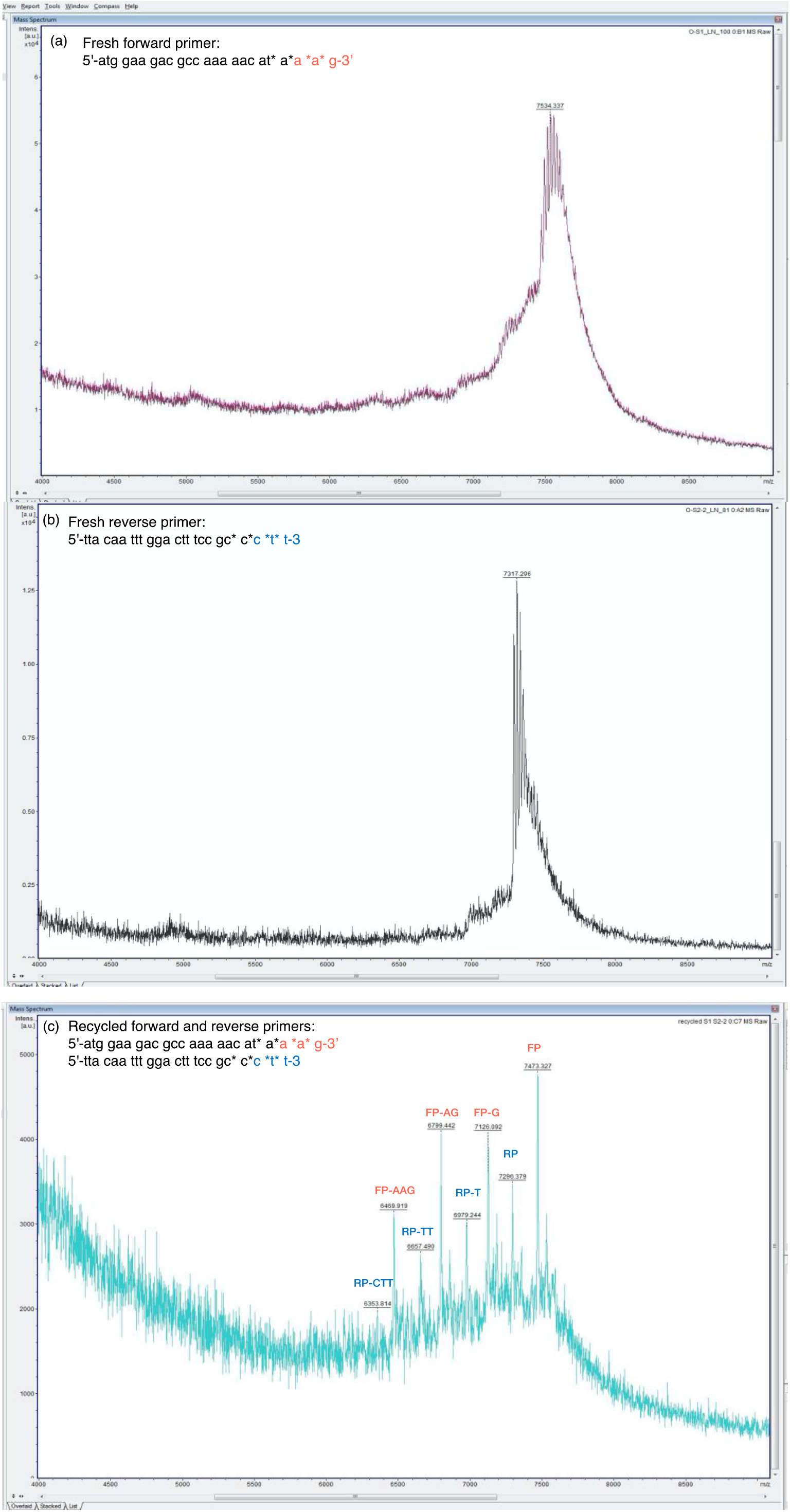
Mass spectrum of (a) 4PS-modified forward primer, (b) 4PS-modified reverse primers, and (c) recycled primers with 1-3 terminal nucleotide lost.

**Figure S5.**
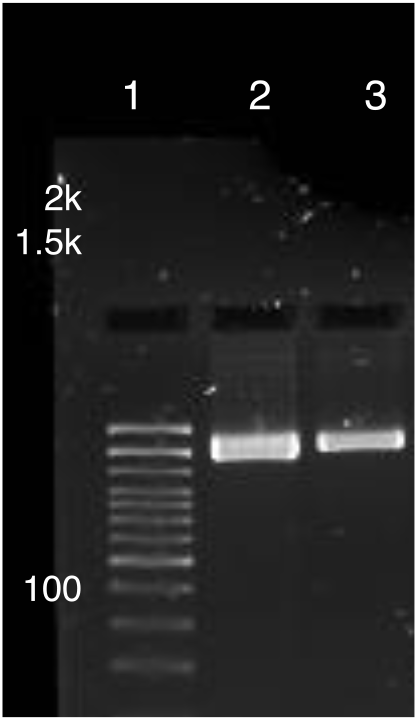
Agarose gel of PCR amplification products from fresh primers (lane 2, 400 nM of added primers) and recycled primers (lane 3, 263 nM of added primers).

**Figure S6.**
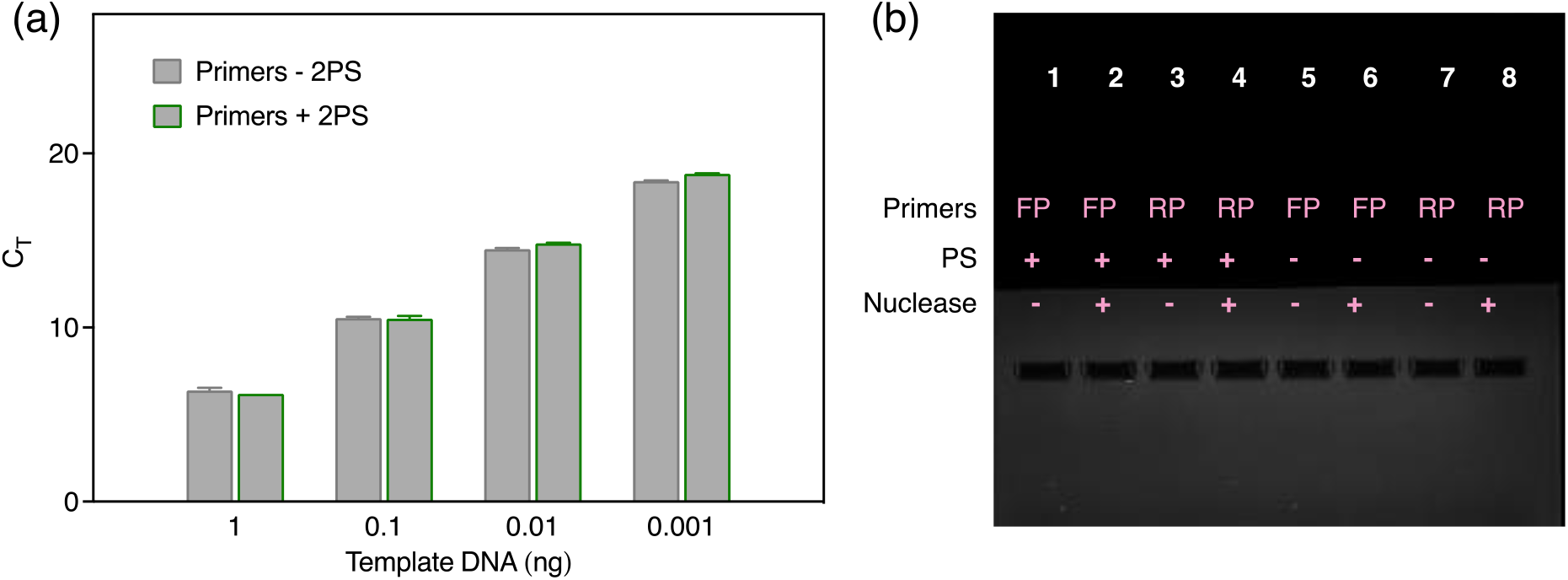
(a) qPCR performance by using primers with or without 2PS-modification with DNA template 1-0.001 ng. (b) nuclease resistance evaluation of primers with or without 2PS-modification. After overnight incubation with nuclease, 10 μL of each primer, or primer + nuclease mixtures were loaded to a 2% agarose gel, lane 1, FP-PS; lane 2, FP-PS + nuclease; lane 3, RP-PS; lane 4, RP-PS + nuclease; lane 5, FP; lane 6, FP + nuclease; lane 7, RP; lane 8, RP + nuclease.

**Figure S7.**
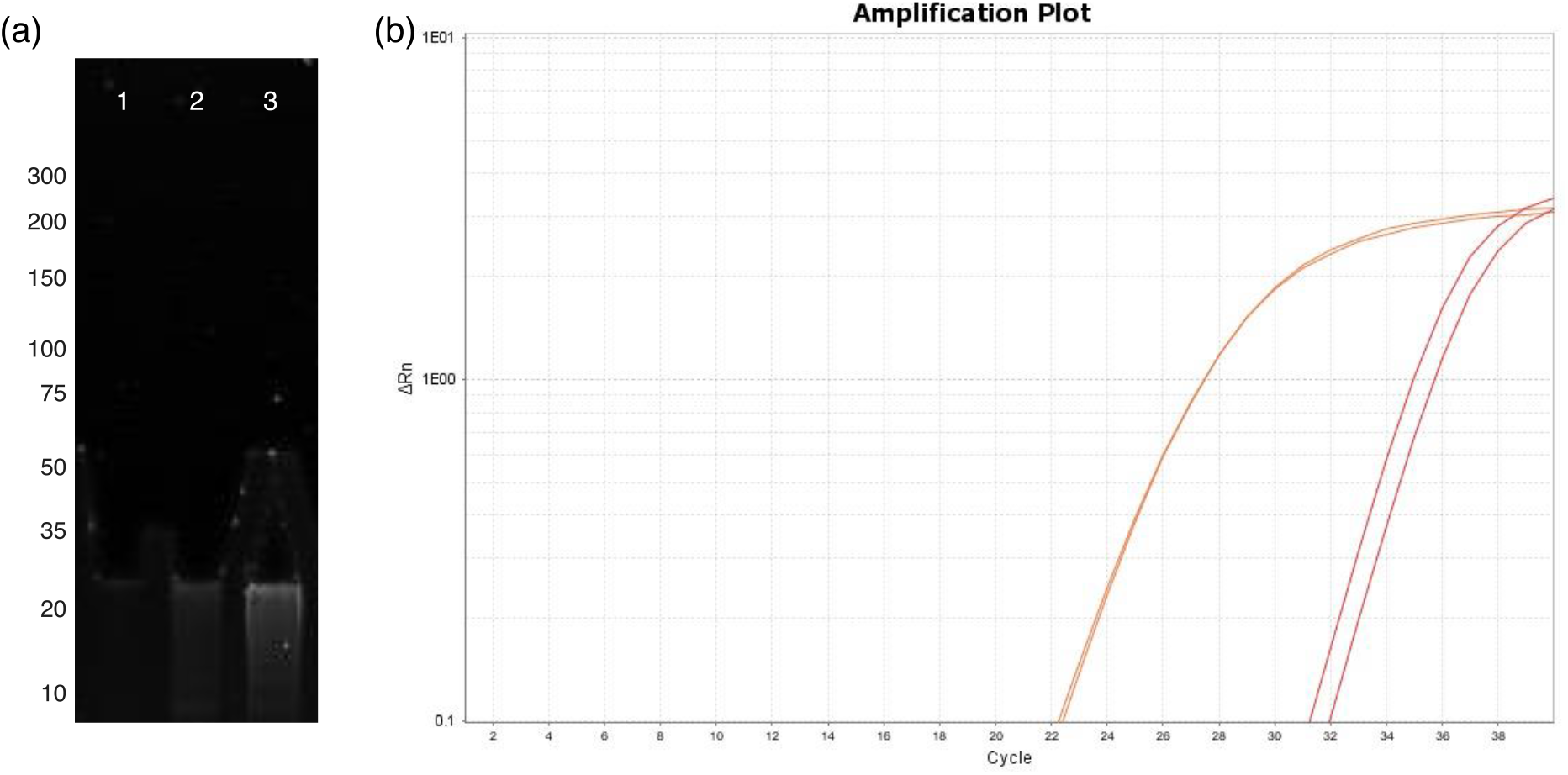
(a) Agarose gel of qPCR amplification product (lane 2) as well as the cleavage fragments (lane 3). (b) qPCR amplification plot of NTC (no template control) sample, red-qPCR kit prepared by recycled primers, orange-qPCR kit prepared by fresh primers.

## REFERENCES

(1) Stahel, W. R. The Circular Economy. Nature 2016, 531 (7595), 435–438. https://doi.org/10.1038/531435a.

(2) Knott, B. C.; Erickson, E.; Allen, M. D.; Gado, J. E.; Graham, R.; Kearns, F. L.; Pardo, I.; Topuzlu, E.; Anderson, J. J.; Austin, H. P.; Dominick, G.; Johnson, C. W.; Rorrer, N. A.; Szostkiewicz, C. J.; Copié, V.; Payne, C. M.; Woodcock, H. L.; Donohoe, B. S.; Beckham, G. T.; McGeehan, J. E. Characterization and Engineering of a Two-Enzyme System for Plastics Depolymerization. Proc. Natl. Acad. Sci. 2020, 117 (41), 25476–25485. https://doi.org/10.1073/pnas.2006753117.

(3) Sathe, D.; Zhou, J.; Chen, H.; Su, H.-W.; Xie, W.; Hsu, T.-G.; Schrage, B. R.; Smith, T.; Ziegler, C. J.; Wang, J. Olefin Metathesis-Based Chemically Recyclable Polymers Enabled by Fused-Ring Monomers. Nat. Chem. 2021, 13 (8), 743–750. https://doi.org/10.1038/s41557-021-00748-5.

(4) Giaveri, S.; Schmitt, A. M.; Roset Julià, L.; Scamarcio, V.; Murello, A.; Cheng, S.; Menin, L.; Ortiz, D.; Patiny, L.; Bolisetty, S.; Mezzenga, R.; Maerkl, S. J.; Stellacci, F. Nature-Inspired Circular-Economy Recycling for Proteins: Proof of Concept. Adv. Mater. 2021, 33 (44), 2104581. https://doi.org/10.1002/adma.202104581.

(5) Liu, W.; Giaveri, S.; Ortiz, D.; Stellacci, F. DNA as a Recyclable Natural Polymer. Adv. Funct. Mater. 2022, 2109538.

(6) Yang, D.; Hartman, M. R.; Derrien, T. L.; Hamada, S.; An, D.; Yancey, K. G.; Cheng, R.; Ma, M.; Luo, D. DNA Materials: Bridging Nanotechnology and Biotechnology. Acc. Chem. Res. 2014, 47 (6), 1902–1911. https://doi.org/10.1021/ar5001082.

(7) Fuller, C. W.; Middendorf, L. R.; Benner, S. A.; Church, G. M.; Harris, T.; Huang, X.; Jovanovich, S. B.; Nelson, J. R.; Schloss, J. A.; Schwartz, D. C.; Vezenov, D. V. The Challenges of Sequencing by Synthesis. Nat. Biotechnol. 2009, 27 (11), 1013–1023. https://doi.org/10.1038/nbt.1585.

(8) Jones, M. R.; Seeman, N. C.; Mirkin, C. A. Programmable Materials and the Nature of the DNA Bond. Science 2015, 347 (6224), 1260901. https://doi.org/10.1126/science.1260901.

(9) Ceze, L.; Nivala, J.; Strauss, K. Molecular Digital Data Storage Using DNA. Nat. Rev. Genet. 2019, 20 (8), 456–466. https://doi.org/10.1038/s41576-019-0125-3.

(10) Li, C.; Faulkner-Jones, A.; Dun, A. R.; Jin, J.; Chen, P.; Xing, Y.; Yang, Z.; Li, Z.; Shu, W.; Liu, D.; Duncan, R. R. Rapid Formation of a Supramolecular Polypeptide–DNA Hydrogel for In Situ Three-Dimensional Multilayer Bioprinting. Angew. Chem. 2015, 127 (13), 4029–4033. https://doi.org/10.1002/ange.201411383.

(11) Udugama, B.; Kadhiresan, P.; Kozlowski, H. N.; Malekjahani, A.; Osborne, M.; Li, V. Y. C.; Chen, H.; Mubareka, S.; Gubbay, J. B.; Chan, W. C. W. Diagnosing COVID-19: The Disease and Tools for Detection. ACS Nano 2020, 14 (4), 3822–3835. https://doi.org/10.1021/acsnano.0c02624.

(12) Hasell, J.; Mathieu, E.; Beltekian, D.; Macdonald, B.; Giattino, C.; Ortiz-Ospina, E.; Roser, M.; Ritchie, H. A Cross-Country Database of COVID-19 Testing. Sci. Data 2020, 7 (1), 345. https://doi.org/10.1038/s41597-020-00688-8.

(13) Sierzchala, A. B.; Dellinger, D. J.; Betley, J. R.; Wyrzykiewicz, T. K.; Yamada, C. M.; Caruthers, M. H. Solid-Phase Oligodeoxynucleotide Synthesis: A Two-Step Cycle Using Peroxy Anion Deprotection. J. Am. Chem. Soc. 2003, 125 (44), 13427–13441. https://doi.org/10.1021/ja030376n.

(14) Palluk, S.; Arlow, D. H.; de Rond, T.; Barthel, S.; Kang, J. S.; Bector, R.; Baghdassarian, H. M.; Truong, A. N.; Kim, P. W.; Singh, A. K.; Hillson, N. J.; Keasling, J. D. De Novo DNA Synthesis Using Polymerase-Nucleotide Conjugates. Nat. Biotechnol. 2018, 36 (7), 645–650. https://doi.org/10.1038/nbt.4173.

(15) Tian, J.; Ma, K.; Saaem, I. Advancing High-Throughput Gene Synthesis Technology. Mol. Biosyst. 2009, 5 (7), 714–722. https://doi.org/10.1039/b822268c.

(16) Praetorius, F.; Kick, B.; Behler, K. L.; Honemann, M. N.; Weuster-Botz, D.; Dietz, H. Biotechnological Mass Production of DNA Origami. Nature 2017, 552 (7683), 84–87. https://doi.org/10.1038/nature24650.

(17) Gibson, D. G.; Young, L.; Chuang, R.-Y.; Venter, J. C.; Hutchison, C. A.; Smith, H. O. Enzymatic Assembly of DNA Molecules up to Several Hundred Kilobases. Nat. Methods 2009, 6 (5), 343–345. https://doi.org/10.1038/nmeth.1318.

(18) Saiki, R.; Gelfand, D.; Stoffel, S.; Scharf, S.; Higuchi, R.; Horn, G.; Mullis, K.; Erlich, H. Primer-Directed Enzymatic Amplification of DNA with a Thermostable DNA Polymerase. Science 1988, 239 (4839), 487. https://doi.org/10.1126/science.239.4839.487.

(19) Liu, W.; Boldt, F.; Tokura, Y.; Wang, T.; Agrawalla, B. K.; Wu, Y.; Weil, T. Encoding Function into Polypeptide-Oligonucleotide Precision Biopolymers. Chem. Commun. 2018, 54 (83), 11797–11800. https://doi.org/10.1039/C8CC04725A.

(20) Roberts, T. C.; Langer, R.; Wood, M. J. A. Advances in Oligonucleotide Drug Delivery. Nat. Rev. Drug Discov. 2020, 19 (10), 673–694. https://doi.org/10.1038/s41573-020-0075-7.

(21) Eckstein, F. Phosphorothioates, Essential Components of Therapeutic Oligonucleotides. Nucleic Acid Ther. 2014, 24 (6), 374–387. https://doi.org/10.1089/nat.2014.0506.

(22) Spitzer, S.; Eckstein, F. Inhibition of Deoxyribonucleases by Phosphorothioate Groups in Oligodeoxyribonucleotides. Nucleic Acids Res. 1988, 16 (24), 11691–11704. https://doi.org/10.1093/nar/16.24.11691.

(23) Wang, L.; Chen, S.; Xu, T.; Taghizadeh, K.; Wishnok, J. S.; Zhou, X.; You, D.; Deng, Z.; Dedon, P. C. Phosphorothioation of DNA in Bacteria by Dnd Genes. Nat. Chem. Biol. 2007, 3 (11), 709–710. https://doi.org/10.1038/nchembio.2007.39.

(24) Fyfe, C.; Sutcliffe, J. A.; Grossman, T. H. Development and Characterization of a Pseudomonas Aeruginosa in Vitro Coupled Transcription-Translation Assay System for Evaluation of Translation Inhibitors. J. Microbiol. Methods 2012, 90 (3), 256–261. https://doi.org/10.1016/j.mimet.2012.05.018.

(25) Yu, S.; Wang, Y.; Li, X.; Yu, F.; Li, W. The Factors Affecting the Reproducibility of Micro-Volume DNA Mass Quantification in Nanodrop 2000 Spectrophotometer. Optik 2017, 145, 555–560. https://doi.org/10.1016/j.ijleo.2017.08.031.

(26) Gilar, M.; Belenky, A.; Budman, Y.; Smisek, D. L.; Cohen, A. S. Study of Phosphorothioate-Modified Oligonucleotide Resistance to 3′-Exonuclease Using Capillary Electrophoresis. J. Chromatogr. B. Biomed. Sci. App. 1998, 714 (1), 13–20. https://doi.org/10.1016/S0378-4347(98)00160-1.

(27) Unlocking P(V): Reagents for chiral phosphorothioate synthesis https://www.science.org/doi/10.1126/science.aau3369 (accessed 2021 -12 -21).

(28) Bustin, S.; Huggett, J. QPCR Primer Design Revisited. Biomol. Detect. Quantif. 2017, 14, 19–28. https://doi.org/10.1016/j.bdq.2017.11.001.

(29) Notomi, T.; Okayama, H.; Masubuchi, H.; Yonekawa, T.; Watanabe, K.; Amino, N.; Hase, T. Loop-Mediated Isothermal Amplification of DNA. Nucleic Acids Res. 2000, 28 (12), E63–E63. https://doi.org/10.1093/nar/28.12.e63.

(30) Shendure, J.; Ji, H. Next-Generation DNA Sequencing. Nat. Biotechnol. 2008, 26 (10), 1135–1145. https://doi.org/10.1038/nbt1486.

(31) C. Nardini; L. Benini; G. De Micheli. Feature - Circuits and Systems for High-Throughput Biology. IEEE Circuits Syst. Mag. 2006, 6 (3), 10–20. https://doi.org/10.1109/MCAS.2006.1688198.

